# High resolution multi-scale profiling of embryonic germ cell-like cells derivation reveals pluripotent state transitions in humans

**DOI:** 10.1101/2025.01.14.632914

**Authors:** Sarah Stucchi, Lessly P. Sepulveda-Rincon, Camille Dion, Gaja Matassa, Alessia Valenti, Cristina Cheroni, Alessandro Vitriolo, Filippo Prazzoli, George Young, Marco Tullio Rigoli, Martina Ciprietti, Benedetta Muda, Zoe Heckhausen, Petra Hajkova, Nicolò Caporale, Giuseppe Testa, Harry G. Leitch

**Affiliations:** Human Technopole, Milan, Italy; Department of Oncology and Hemato-Oncology, University of Milan, Milan, Italy; MRC Laboratory of Medical Sciences (LMS), London, UK; Institute of Clinical Sciences (ICS), Faculty of Medicine, Imperial College London, London, UK; Department of Experimental Oncology, European Institute of Oncology IRCCS, Milan, Italy; Laboratory of Endometrium, Endometriosis and Reproductive Medicine, Department of Development and Regeneration, KU Leuven, Leuven, Vlaams-Brabant, Belgium and Laboratory of Ion Channel Research, Department of Molecular Medicine, KU Leuven, Leuven, Belgium; Department of Medical Biotechnology and Translational Medicine, University of Milan, Milan, Italy; UCL Great Ormond Street Institute of Child Health, Genetics & Genomic Medicine Department, London, UK and North East Thames Regional Genetic Service, Great Ormond Street Hospital for Children NHS Foundation Trust, London UK

## Abstract

Primordial germ cells (PGCs) are the embryonic precursors of the gametes. In mice and rats, PGCs can readily acquire pluripotency in vitro by forming embryonic germ cells (EGCs). To date, a comparable in vitro system has not been established in humans, despite the fact that human PGCs (hPGCs) readily undergo pluripotent conversion in the context of germ cell tumorigenesis. Here we report that hPGC-like cells (hPGCLCs) undergo conversion to human embryonic germ-like cells (hEGCLCs) upon exposure to the same inductive signals previously used to derive mouse EGCs. This defined, feeder-free culture system allows efficient derivation of human EGCLCs which can be expanded and maintained in standard human pluripotent stem cell medium. hEGCLCs are transcriptionally similar to human pluripotent stem cells (hPSCs) and can differentiate into all three germ layers, as well as giving rise to PGCLCs once more - demonstrating the interconvertibility of pluripotent states. This is also evident at the epigenetic level, as the initial DNA demethylation that occurs in hPGCLCs is largely reversed in hEGCLCs, restoring DNA methylation to the level observed in hPSCs. This new in vitro model captures the transition from the pluripotent stem cell state to a germ cell identity and back again, and therefore represents a highly tractable system to study pluripotent and epigenetic transitions, including those which occur during human germ cell tumorigenesis.

**Graphical abstract:** 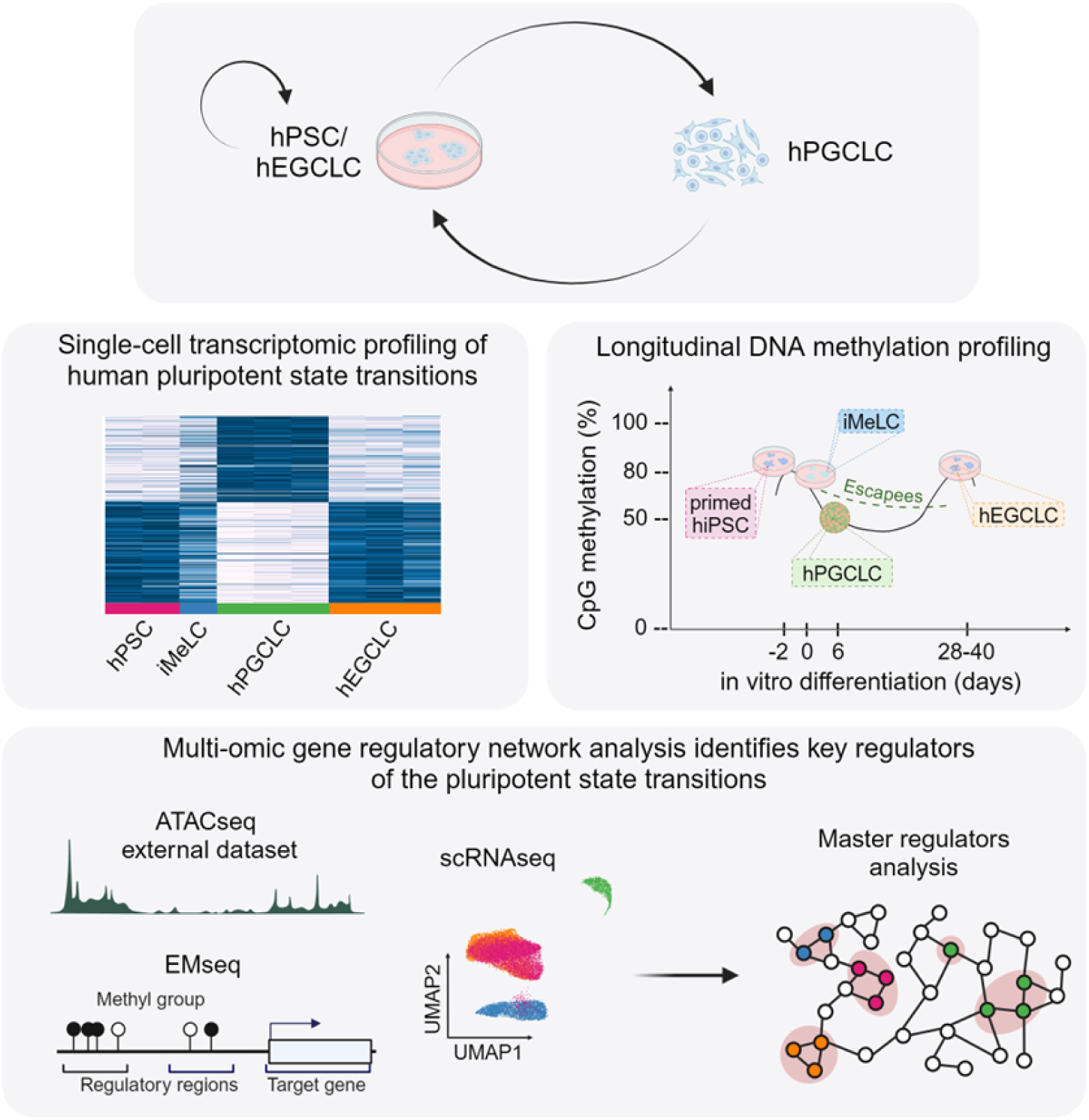

**In brief:** We report the first fully defined system to efficiently convert hPGCLCs to a pluripotent stem cell (PSC) state. We tracked pluripotent state transitions by multi-omic analysis and provided a high-resolution map of the transcriptional and epigenomic transitions upon entry to and exit from the human germline.

**Highlights:**

- Efficient derivation of hEGCLC in fully defined feeder-free conditions
- Single-cell transcriptomic profiling of transitions from the hPSC state to hPGCLCs and back.
- Longitudinal DNA methylation profiling highlights the overall reversibility of epigenetic states
- Multi-omic gene regulatory network analysis identifies key regulators of pluripotent transitions

## Introduction

In sexually reproducing organisms, germ cells play a crucial role in transmitting the genetic and epigenetic information necessary for the development of a new organism, serving as an enduring link between generations ^1,2^. Primordial germ cells (PGCs) are the founder cells of the germ cell lineage and as such the earliest embryonic precursors of the gametes ^3^. In humans, PGC specification likely occurs around embryonic day 12–16 (post-conceptional week 2), either in the posterior epiblast of the early post-implantation embryo or the nascent amnion or both sites ^4^. At week 3–5, human PGCs (hPGCs) migrate from the yolk sac wall through the hindgut and colonise the developing gonads, where sex-specific differentiation occurs from around week 9 ^5–85–85,8,9^During their migratory phase, hPGCs commence a wave of global DNA demethylation, including significant loss of DNA methylation at imprinting control regions ^5–7^, which heralds the most extensive epigenetic reprogramming process during development. In addition to erasure of imprints, PGC reprogramming may function to enable the activation of the gametic and meiotic programs, enable X chromosome reactivation, erase epimutations, or regulate retrotransposons^10,11^.

PGCs are traditionally regarded as unipotent, as physiologically they give rise only to gametes. However, mouse PGC development is dependent on the expression of pluripotency genes ^12–14^ and PGCs exhibit pluripotent characteristics. For instance, PGCs are the cell of origin of pluripotent germ cell tumours such as teratocarcinomas, from which mouse embryonal carcinoma (EC) cells were first derived ^15,16^. In addition, when placed in culture, mouse PGCs can directly give rise to pluripotent stem cells called embryonic germ (EG) cells ^17,18^. Mouse EG cells can contribute to chimaeras ^19,20^ and their properties are indistinguishable from naive embryonic stem (ES) cells, despite their different origin ^21^. In fact, mouse EG cell conversion from PGCs is highly efficient, calling into question the designation of PGCs as a unipotent lineage ^22^. Indeed, we have recently revisited the classic chimaera experiments which investigated the potency of mouse PGCs ^23^. Our findings suggest that PGCs exhibit a latent form of pluripotency, that distinguishes them from somatic precursor cells ^24^.

The latent pluripotency of PGCs is not a mouse-specific property. Naive pluripotent EG cell lines have also been derived from rats ^25–27^, completing the naive pluripotent stem cell repertoire in this species, following the derivation of rat ES cells ^28,29^ and induced pluripotent stem cells (iPSCs) ^30^. In humans, hPGCs can give rise to germ cell tumours, many of which have pluripotent properties^31^. Notably, 50% of paediatric germ cell tumours occur outside the gonad, anywhere from coccyx to cranium, originating from mismigrated hPGCs ^32^. However, although derivation of human ES cells ^33^ and iPSCs ^34^ were reported decades ago, to date human EG cells that exhibit the same properties have not been derived ^35^. Despite the fact that hPGCs can give rise to pluripotent tumours, it has been suggested that species-specific differences - such as the lack of expression of SOX2 in hPGCs - might explain the inability to derive human EG cells ^36^. Further progress towards the derivation of human EG cells has also been hampered by the lack of access to hPGCs, in particular at the pre-migratory and migratory stages which are known to be most amenable to EG cell conversion in mice ^20,22^.

The recent advances in in vitro induction of human PGC-like cells (hPGCLCs) has provided a new and exciting tool to study the early human germline ^37,38^. Of note, a number of labs have developed culture conditions for hPGCLCs which allows expansion of hPGCLC numbers while maintaining their PGC-like properties ^39–41^. Modifications to the media composition can allow the emergence of pluripotent stem cell colonies from proliferating cultures of hPGCLCs ^41^. However, a defined culture system to trigger the conversion of hPGCLCs to pluripotent stem cell lines has not been reported. We previously reported the first fully defined, feeder-free culture system for the derivation of mouse EG cells ^22,42^. Here we report the development of a fully-defined, feeder-free culture system that enables the transition of human germline cells (hPGCLCs) to pluripotent stem cells, called human embryonic germ cell-like cells (hEGCLCs). We demonstrate the utility of our novel culture system by undertaking high resolution, multi-omic profiling during the conversion of hPSCs to hPGCLCs and back again, producing a rich dataset that provides insight into these unique and clinically relevant pluripotent transitions, and providing a comprehensive resource for the broader research community.

## Results

### hEGCLCs derivation from hPGCLCs in defined and feeder-free conditions

Given the similar requirements for PGCLC induction between mouse and human, we hypothesised that the factors that trigger conversion of PGCs (or PGCLCs) to pluripotent stem cells might also be conserved. First, hPGCLCs were differentiated from hPSCs through incipient mesoderm like-cells (iMeLCs), using a well-established protocol ^38^ (Figure 1A, 1B). At day 4 or 6, hPGCLCs were identified by the expression of BLIMP1 and TFAP2C fluorescent reporters (BTAG) ^38^ or by using previously published cell surface markers, EpCAM (CD326) and INTEGRINα6 (CD49f) ^38,43^ (Figure 1C). After sorting, hPGCLCs were plated on geltrex in an induction medium containing the same factors that trigger efficient EG cell conversion in mice (basal medium supplemented with human leukemia inhibitory factor (hLIF), the GSK3 inhibitor CHIR99021 (CH), Forskolin (FK), basic fibroblast growth factor (bFGF), human stem cell factor (hSCF) and retinoic acid (RA) (see Methods)). After 48 hours, the medium was transitioned to standard human pluripotent stem cell media by half medium changes. In these defined and feeder-free culture conditions we observed the emergence of colonies that resembled human pluripotent stem cells. After 14 days these colonies were picked, expanded and maintained using standard human pluripotent stem cell culture protocols (Figure 1A, 1B). Using this protocol we have derived multiple such cell lines from hPGCLCs themselves induced from both hiPSCs and ESCs (male and female). In keeping with a previous study ^41^, we have designated these cell lines as hEGCLCs to reflect their origin from hPGCLCs.

**Figure 1.**
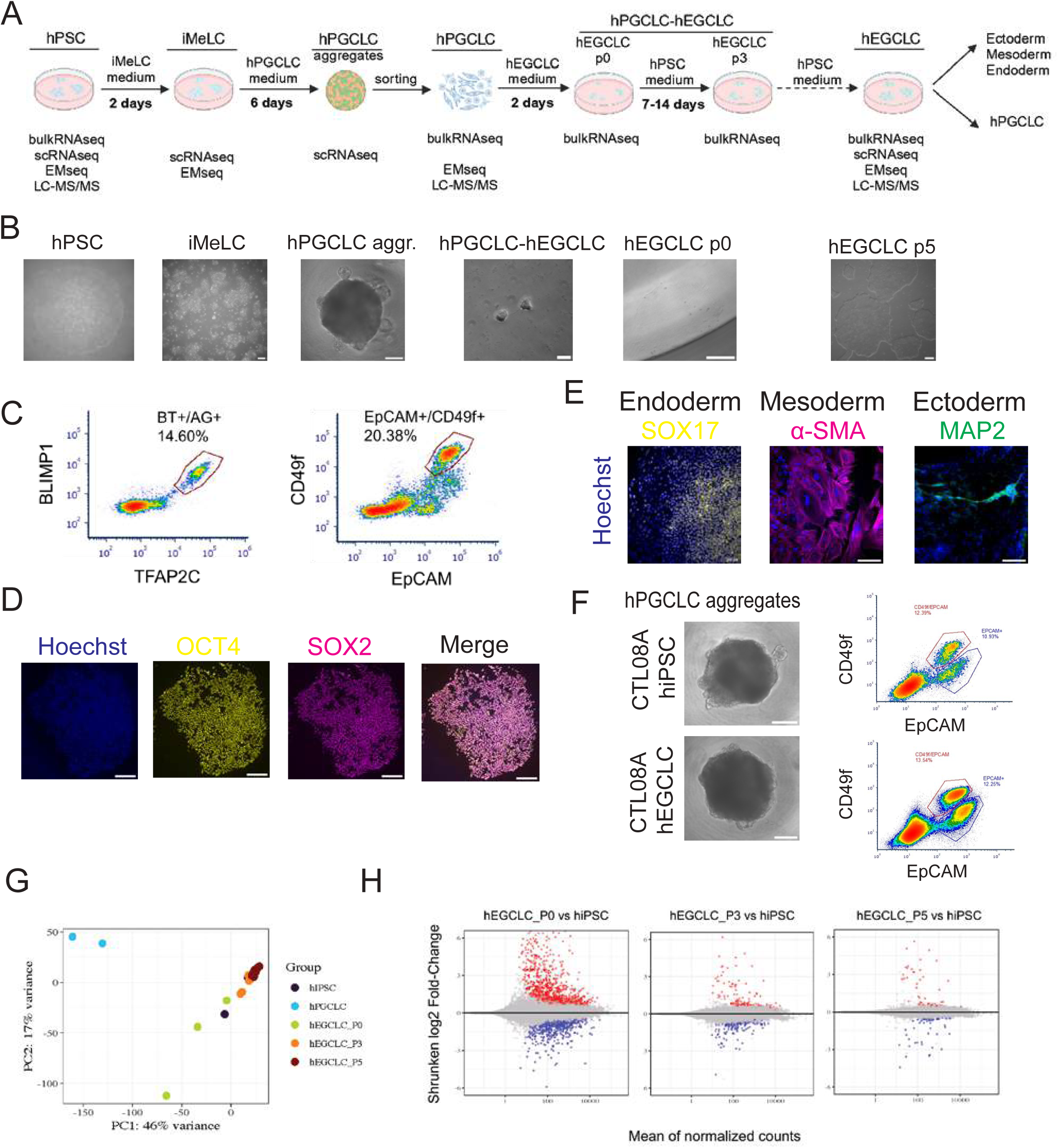
hEGCLCs derivation from hPGCLCs in defined and feeder-free conditions and validation of their pluripotency. **(A)** Steps to differentiate hPGCLCs from hPSCs and reprogram them into hEGCLCs and schematic of omics assays performed. hPSCs are differentiated in 2 days into iMeLCs, and further into hPGCLCs, within 3D aggregates. hPGCLCs aggregates are dissociated and hPGCLCs sorted (10 cells for each well of 96 multiwell) and grown for 2 days in a hEGCLC induction medium, containing hLIF, CHIR, forskolin, bFGF, hSCF and Retinoic Acid. After 2 days, hEGCLC induction medium is gradually switched into conventional hPSCs culture media, allowing the reprogramming of hPGCLCs into hEGCLCs. hEGCLCs are able to differentiate into the three germ layers (ectoderm, mesoderm, endoderm) and into hPGCLC **(B)** Phase contrast images exemplifying the key steps and cellular states of the differentiation protocol (scale bar: 250μm; 25μm for hPGCLC-hEGCLC image) **(C)** Representative flow-cytometry density plot showing the percentage of hPGCLCs within hPGCLC aggregates (double positive cells for: BLIMP-tdTomato and AP2γ-GFP, BTAG reporter line, left panel; EpCAM-FITC and CD49f-PE, right panel) **(D)** Representative immunofluorescence images of hEGCLCs colonies (passage 5) stained for pluripotency markers: SOX2, OCT4; nuclei stained with Hoechst (scale bar: 150μm) **(E)** hEGCLC tri-germ layer differentiation: representative immunofluorescence staining for lineage specification markers: SOX17 (endoderm), α-SMA (mesoderm), MAP2 (ectoderm); nuclei stained with Hoechst (scale bar: 150 μm) **(F)** On the left: representative phase contrast images of day6 hPGCLC aggregates differentiated in parallel from CTL08A hiPSCs (top panel) and hEGCLCs (bottom panel); scale bar 250 μm. On the right: corresponding flow cytometry density plots of CD49f (Integrin-α6, Y-axis) and EpCAM (X-axis) surface markers. Percentages of double-positive cells (i.e. hPGCLCs) are shown in the red gate **(G)** Principal Component Analysis (PCA) of bulk RNASeq profiling of the samples indicated in the legend (hiPSCs, day 4 hPGCLCs, hEGCLCs passage zero [P0, 14 days after EGC induction], hEGCLCs P3 and hEGCLC P5) **(H)** MA-plot showing the mean normalized counts (X-axis) and shrunken log2Fold-Change values of expressed genes, for the following pair-wise comparison: hEGCLC P0 vs hiPSC; hEGCLC P3 vs hiPSC; hEGCLC P5 vs hiPSC (from left to right respectively). Differentially expressed genes (DEGs) after p-value adjustment (q-value<0.01) are displayed in red (upregulated) or in blue (downregulated)

Next, we monitored the transition from hPGCLC-to-hEGCLC using live-cell imaging. Cultured hPGCLCs initially exhibited a typical migratory morphology, but over the first week of culture small colonies began to form and by day 14 colonies with the typical appearance of human pluripotent stem cells were present (Figure 1B). Live imaging of the BTAG fluorescent reporters ^38^ indicated that the hPGC hallmark genes TFAP2C and BLIMP1 are downregulated from around day 7 of culture and are no longer expressed in hEGCLC lines (Video S1 and S2). In addition, we performed live-imaging using a previously published SOX2 reporter cell line ^44^, which revealed a reciprocal transcriptional upregulation of the pluripotency gene *SOX2* over a similar time frame. As expected, SOX2 expression was maintained thereafter in hEGCLCs (Video S3). Thus, these live-imaging experiments indicate that downregulation of the PGC programme occurs in a similar timeframe to the upregulation of SOX2, and also rule out that hEGCLC colonies emerge from contaminating undifferentiated pluripotent stem cells. hEGCLCs can be expanded for more than 30 passages using conventional pluripotent stem cell culture conditions, maintaining karyotypic stability. Finally, plating 10 hPGCLCs per well of a 96 well plate, we counted the number of hEGCLC colonies that emerged after 14 days. This revealed a derivation efficiency of up to 10-15% (Figure S1A), which approaches the highest efficiency reported for mouse EG cell derivation from wild-type PGCs ^22^.

At the protein level, hEGCLCs express pluripotency markers SOX2, OCT4, NANOG and SSEA-4 and no longer express PGC markers such as TFAP2C, BLIMP1 and SOX17 (Figure 1D, S1B, and S1C). We also tested the differentiation capacity of hEGCLCs into the three germ layers, confirming their pluripotency, by using standard embryoid body formation assays ^45,46^ (Figure 1E, S1D). Additionally, hEGCLCs are able to differentiate into hPGCLCs, with an efficiency comparable to that of the hiPSC parent line (Figure 1F). Thus, hEGCLCs show the broad in vitro differentiation capacity expected from pluripotent stem cell lines. These results establish the first fully-defined and highly efficient method to derive hEGCLCs.

### Transcriptional profiling of the hPGCLC-to-hEGCLC transition

A benefit of an efficient and defined cell culture system is that it facilitates downstream molecular profiling. We first undertook transcriptomic profiling by bulk RNA-sequencing of the hiPSC parental line, day 4 hPGCLCs, and hEGCLCs at passage 0 (P0, 14 days after hEGCLC induction), P3 (hEGCLCs P3) and P5 (hEGCLCs P5) (Figure 1G). This revealed that by passage 5, hEGCLCs cluster together with hiPSCs in principal component analysis (PCA) pointing to the similarity of their transcriptomes (Figure 1G). Further, the number of differentially expressed genes (DEGs) between hEGCLCs and hiPSCs is reduced from 467 at P0 to only 46 at P3 and 50 at P5 (Figure 1H and Table S1). The high number of differentially expressed genes at P0 likely reflects a still mixed population of hEGCLCs and non-reprogrammed cell types. Established hEGCLC lines express pluripotency genes at similar levels to hiPSCs and have downregulated PGC genes (Figure S1E). These results confirm that hEGCLCs established from hPGCLCs are transcriptionally similar to hiPSCs, further confirming their identity as pluripotent stem cells.

To increase the resolution of transcriptome analysis we profiled hiPSC, iMeLC, day 6 hPGCLC aggregates and hEGCLC transcriptomes at the single-cell level, obtaining 27925 high quality cells after standard quality checks and filters (see Methods). To reduce the dimensionality of the dataset, we applied principal component analysis (PCA, Figure S2A) on highly variable genes, selected with Triku ^47^ (see Methods) and uniform manifold approximation and projection (UMAP) (Figure S2B).

We first confirmed hiPSC, hEGCLC and iMeLC identity by checking the expression of pluripotency (SOX2, DPPA4, PRDM14, ZIC5) and iMeLC (EOMES, SP5, MIXL1) markers, respectively (Figure 2A left panel, S2C and D). This first instance of hEGCLC scRNAseq profiling exposed their remarkable, hiPSC-equivalent degree of homogeneity ^48^ (Figure 2A left panel and S2B). We then systematically annotated the identity of each cell population captured in our in vitro developmental transitions by analysing the expression of validated markers from the existing knowledge bases ^6,38,49–53^, including markers from a reference single-cell dataset of a gastrulating human embryo ^54^ (Figure 2A left panel and Table S2). We annotated cells expressing PGC markers (NANOS3, CD38, KLF4, TFAP2C, PRDM1, SOX17, KLF4, PIFO) as hPGCLCs; cells expressing amnion markers (TFAP2A, GATA3, ISL1, WNT6, GABRP, HAND1) as amnion-like cells (AmLCs), cells expressing endoderm markers (FOXA2, APOA1, APOB, TTR, FGB, MTTP) as endoderm-like cells (EndLCs), cells expressing hemato-endothelial markers (MEF2C, GMFG, LAPTM5, ICAM2, LMO2) as hemato-endothelial progenitors (HEPs) and cells expressing mesoderm markers (HAND1, GATA6, FOXF1, SNAI2) as mesoderm-like cells (MLCs) (Figure 2A left panel, S2C and SD). These data outline a heterogeneous composition of hPGCLC aggregates (Figure S2B, green cluster), recapitulating cell types present in the early phases of human embryo development ^55,56^, as also highlighted in other studies ^57,58^.

**Figure 2.**
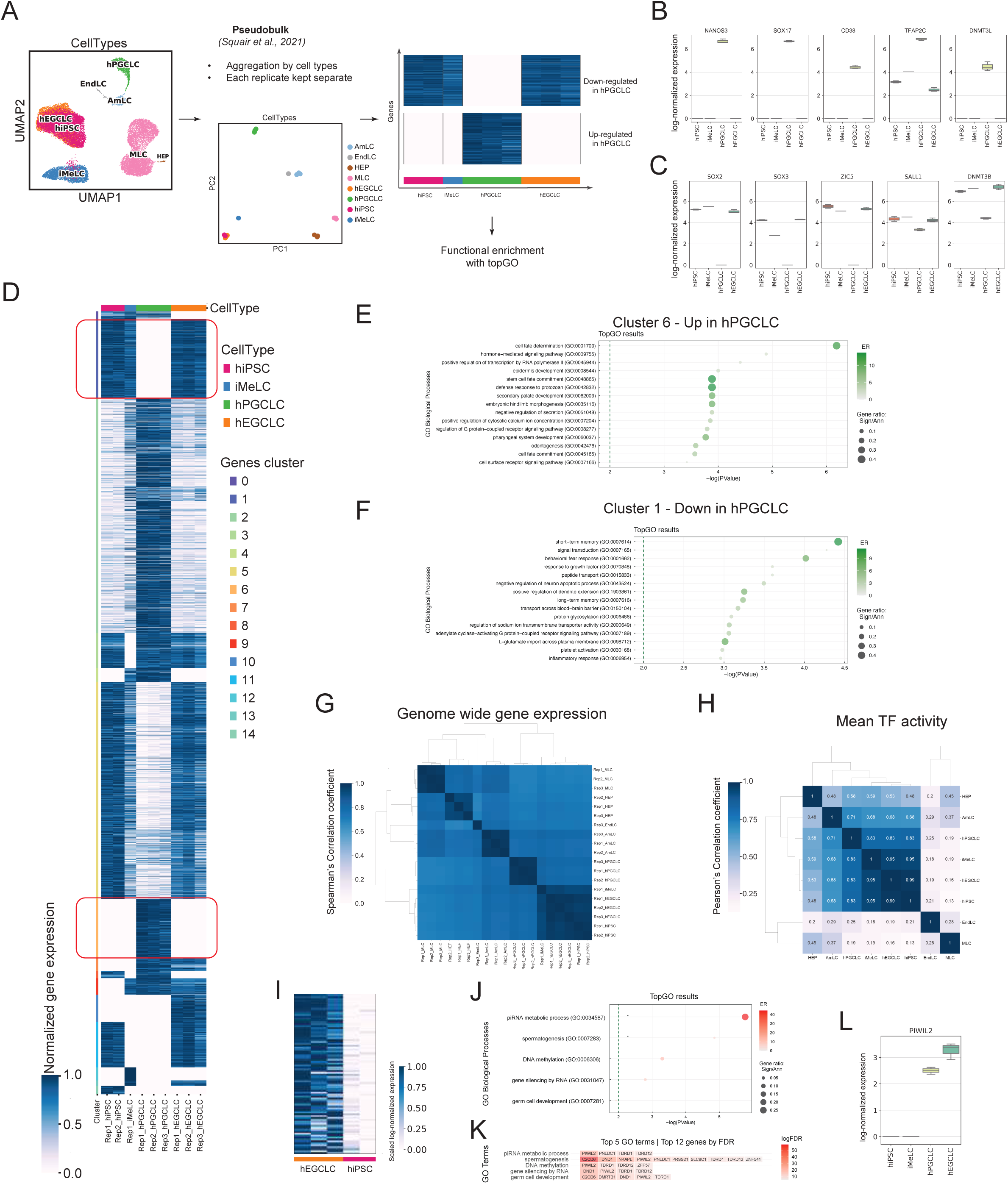
In vitro model longitudinal transcriptomic profiling at single-cell resolution **(A)** Schematic of the pseudo-bulk differential expression analysis done on the single-cellRNAseq data. From left to right: UMAP visualization of all cells in the dataset, after pre-processing and filtering, colored by annotated cell type (hiPSC =induced Pluripotent Stem Cell; iMeLC = incipient Mesoderm-Like Cells; hPGCLC = human Primordial Germ Cell-Like Cells; hEGCLC = Embryonic Germ Cell-Like Cells; MLC = Mesoderm-Like Cells; AmLC = Amnion-like cells; EndLC = Endoderm-like cells; HEP = Haemato-Endothelial Progenitors). PCA plot of pseudo-bulk counts, colored by annotated cell types. Unsupervised approach to identify clusters of gene expression patterns, then used as input for functional enrichment analysis. **(B)** and **(C)** Box plots of pseudo-bulk log-normalized level of expression (Y-axis) of representative genes found up or down-regulated in hPGCLC, compared to the other three key cell types of our dataset (ordered longitudinally on the X-axis: hiPSC, iMeLC, hEGCLC). **(D)** Heatmap of genes clusters obtained as described in (A) Clusters of genes up-regulated (Cluster 6) or down-regulated (Cluster 1) in hPGCLC compared to the other three key cell types are highlighted with a red box. **(E)** and **(F)** Dot plot of significantly enriched Gene Ontology (GO) terms (Biological Processes category, BP) for gene cluster 6 (E) and 1 (F), highlighted in (D) Y-axis: GO term; X-axis: -log(p-value)>2. Color gradient indicates the enrichment ratio (ER) > 2, dot dimension indicates gene ratio. **(G)** Clustered heatmap of Spearman correlation coefficient between annotated cell types. **(H)** Heatmap of Pearson correlation coefficient of TF activity between annotated cell types. **(I)** Heatmap of pseudo-bulk gene expression of the 82 genes upregulated in hEGCLC compared to hiPSC. Color gradient: scaled log-normalized expression. **(J)** Dot plot of significantly enriched Gene Ontology (GO) terms (BP category) of DEGs in **(I)** Y-axis: GO terms; X-axis: -log(p-value)>2. Color gradient indicates the enrichment ratio (ER) > 2, dot dimension indicates gene ratio. **(K)** Top 12 genes by false discovery rate (FDR) for each of the top 15 GO terms shown in panel (J) Color gradient indicates logFDR. **(L)** Box plots of pseudo-bulk log-normalized level of expression (Y-axis) of PIWIL2 gene in hiPSC, iMeLC, hPGCLC and hEGCLC (X-axis).

To compare our in vitro model with publicly available in vitro data, we projected our longitudinal scRNAseq data on the scRNAseq dataset from an external in vitro reference ^59^, where hPGCLCs were differentiated from human embryonic stem cells (hESCs) using the same protocol ^38^ (Figure S2E). We confirmed how our hPGCLCs map onto the published NANOS3^+^ cells (i.e. hPGCLCs), as well as iMeLCs onto iMeLCs and hiPSCs onto hESCs (Figure S2F). Interestingly, hEGCLCs, present only in our dataset, map onto published hESCs, further confirming their transcriptional overlap with hPSCs (Figure S2F), as also evident from their quite even distribution among the clusters we annotated as hEGCLC and hiPSC in our dataset (Figure 2A left panel, S2F and Table S2).

Finally, we benchmarked our in vitro data with in vivo fetal references (human prenatal gonads from 6–16 weeks post-fertilization ^60,61^, Figure S2G) noting how our hPGCLCs differentiated in vitro resemble the in vivo fetal counterpart (Figure S2H).

### Longitudinal gene expression patterns reveal the emergence and dissolution of germline identity

To identify gene expression patterns throughout differentiation, we analysed our scRNAseq data through a pseudo-bulk approach ^62–64^ to leverage the statistical rigour of generalised linear models for differential expression analysis ^65^. We aggregated cells according to the annotated cell types (see above, Figure 2A), keeping each replicate separated and found that different replicates of the same cell type cluster together, as do hiPSCs and hEGCLCs (Figure 2A middle panel). This is also confirmed by Spearman’s correlation analysis (Figure 2G), underlining the degree of consistency across replicates and, once again, the hiPSC-hEGCLC transcriptional overlap.

Next, we defined groups of DEGs showing specific longitudinal expression patterns, in the four key cell types of our in vitro model (namely hiPSCs, iMeLCs, hPGCLCs and hEGCLCs), using an unsupervised approach. We computed the log fold-change (logFC) of genes in the latter three cell types with respect to their expression levels in hiPSC, filtered by a minimum significance of 0.01 and then used the logFC to identify groups of interest ^66^ (Methods, Figure 2A right panel). We identified 15 clusters (Table S3), as shown in the heatmap in Figure 2D. Of particular interest are cluster 2 and 6 that include genes (such as NANOS3, SOX17, CD38, TFAP2C and DNMT3L, Figure 2B) whose expression along differentiation from hiPSC to hEGCLCs increases in hPGCLCs and decreases again in hEGCLCs and cluster 5 and 1 that, conversely, include genes (such as SOX2, SOX3, ZIC5, SALL1 and DNMT3B, Figure 2C) whose expression decreases in hPGCLCs and increases again in hEGCLCs. These expression patterns are aligned with what is already known from hiPSC-to-hPGCLC developmental transition ^38,59^, serving as positive controls that confirm the robustness of the experimental and analytical approach and adding new insights into hPGCLC-to-hEGCLC transition. We then performed functional enrichment analysis ^67^ on these DEGs, focusing on cluster 6 and 1, which show the strongest up/down-regulation in hPGCLCs compared to the other three cell types. For cluster 6 (genes upregulated in hPGCLCs), we identified terms related to cell fate determination and embryo development and morphogenesis (Figure 2E). Interestingly, top genes (by FDR) associated with these terms are SOX17, CD38, ITGB3 and GATA transcription factors, known early primordial germ cells markers ^37,68^, genes related to WNT signalling pathway (such as WNT7A ^58,69^), already known to be upregulated upon hPGCLC differentiation ^49,70,71^ as well as genes involved in PGC migration (such as EDN1, MSX1 and MSX2 ^49,68,72^) (Figure S2I). Instead, among DEGs from cluster 1 (genes downregulated in hPGCLCs), we identified terms related to differentiation towards other lineages, such as regulation of dendrite extension, platelet activation and inflammatory response (Figure 2F and S2L). This is consistent with the need to not only activate germ-cell specific genes but also to repress genes specific for other cell lineages ^38,49,73,74^.

Next, to investigate the regulatory logic of the developmental transitions exposed by our longitudinal design, we performed transcription factor (TF) activity inference using decoupleR package ^75^ and DoRothEA regulons collection ^76^ (see Methods). First, we conducted a correlation analysis of TF activity that highlighted the highest regulatory similarity between hiPSCs and hEGCLCs (Pearson’s correlation coefficient equal to 0.99) (Figure 2H). Among the 292 TFs considered in the analysis (see Methods), we selected those relevant to pluripotency, germ cell and endoderm specification pathways and plotted them in a UMAP to visualise the inferred TF activity scores. NANOG and OCT4 (i.e. POU5F1) scores are higher in hiPSC and hEGCLC clusters, whereas TFAP2C score is higher in AmLC and hPGCLC clusters and FOXA2 score in EndLC cluster, consistent with our own and others’ previous findings ^58^ (Figure S2M).

To deepen the investigation of finer differences between hiPSCs and hEGCLCs, we performed differential expression analysis focusing specifically on this comparison. We used the pseudobulk approach and found 34 genes downregulated (Figure S2N and Table S3) and 82 genes upregulated (Figure 2I and Table S3) in hEGCLCs compared to hiPSCs. By performing a functional enrichment analysis on the hEGCLC upregulated genes, we found terms related to piRNA metabolic processes and spermatogenesis, with PIWIL2 as one of the top genes associated with these terms ^77^ (Figure 2J, 2K). Since its expression level is higher in hPGCLCs than in hiPSCs and then does not decrease in hEGCLCs (Figure 2L), we speculate that it could disclose a molecular memory of their origin from hPGCLCs.

In summary, this longitudinal transcriptomic profiling at single-cell resolution elucidated the gene expression patterns and the transcriptional logic underlying the transitions from the hPSC state to hPGCLCs and back.

### Gene regulatory network analysis highlights key regulators of hEGCLC transcriptional program

To better investigate the key transcription factors and regulatory logic governing the reprogramming from PSCs to germ cells and back, we performed gene regulatory network (GRN) analysis using CellOracle ^78^,leveraging our own scRNAseq dataset alongside publicly available ATACseq profiles ^79,80^. For GRN construction, we associated each regulatory region (identified by chromatin accessibility) to a target gene (among the HVGs and a list of selected genes, see Methods and Table S4) and to one or more TFs, building TF-target gene(s) graphs. From this first GRN, we harnessed our transcriptomic data to build four different cell-type specific GRNs (hiPSC, iMeLC, hPGCLC, hEGCLC GRN, Table S4).

First we considered two key GRN parameters, the betweenness centrality (i.e. a measure proportional to the number of times each TF connects to other nodes in the network, that gives an idea of the importance of each TF in that GRN, Figure 3A) and the out-degree centrality (i.e. a measure of the number of edges that go out from that node, a measure of the number of targets regulated by that TF within the network, Figure S3A). This analysis revealed that the top regulators of hiPSC and hEGCLC GRN are MYC, HDAC2, ZIC5 and SP3. Instead for iMeLC, EOMES and SP5 emerged as master regulators, as expected ^38^, together with TFs such as ENO1 and HDAC2 that are in common with their pluripotent precursors. As for hPGCLC, among the top regulators we identified TFAP2C, PRDM1 (i.e. BLIMP1), and KLF4, in line with what is already known in literature ^3,49,68,79^, confirming the robustness of our analysis, but also REST that has been only recently identified as an important TF in PGCs ^81^ (Figure 3A and S3A). These top regulators emerged also when comparing GRNs of these cell types in pairs (Figure 3B).

**Figure 3.**
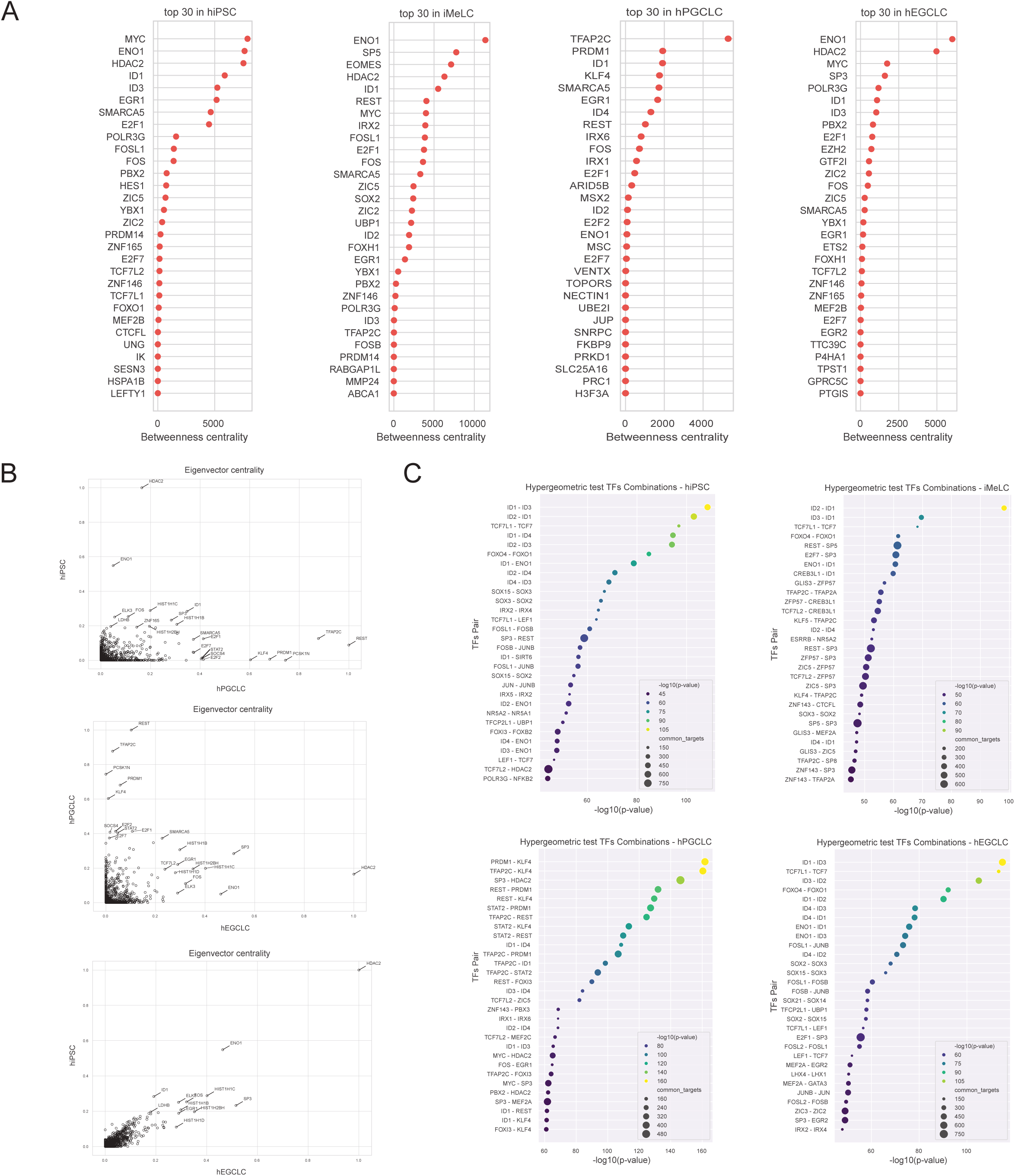
Gene regulatory network analysis highlights key regulators of hEGCLC transcriptional program **(A)** Dot plot of top-ranking transcription factors (TF) based on betweenness centrality (X-axis) measured for each cell type (reported in each plot title). **(B)** Scatter plot of eigenvector centrality measured for each TF in each cell type. Each plot represents the comparison of two cell types (reported in the axis title). hiPSC vs hPGCLC, hPGCLC vs hECGLC and hiPSC vs hEGCLC are reported (y vs x-axis) from top to bottom, respectively. **(C)** Dot plot of enrichments measured in each cell-type for the overlap of targets for each couple of TFs considered. Top 30 TF couples are reported based on significance [-log10(p-value) of the hypergeometric test measured on the overlap between targets of the two TF reported on the Y-axis, with respect to the genes expressed in each cell type]. Dot size is proportional to the size of the intersection. Color is proportional to significance. hiPSC and iMeLC top TF couples are reported from left to right on top of the panel. hPGCLC and hEGCLC top TF couples are reported from left to right on the bottom of the panel.

We then assessed the cooperativity among the TFs of the GRN, for each cell type, by using a hypergeometric test to evaluate the statistical significance of their overlap in terms of shared targets. In hiPSC and hEGCLC GRNs, among the top cooperating TFs we identified TF belonging to ID, TCF, FOXO and SOX families, known to be associated with pluripotency ^82–85;^ in iMeLC again ID, TCF, FOXO, but also REST, SP3 and SP5; in hPGCLC KLF4, PRDM1, TFAP2C, REST and STAT2 (Figure 3C). We also performed the same analysis, in a supervised way, focusing on known TFs of interest, confirming the results obtained so far and highlighting the high number of common targets between top cooperating TFs for each cell type. For example, the pairs MYC-HDAC2 and MYC-SP3 in hiPSC and hEGCLC with 761 and 811, 742 and 806 common targets, respectively, and by TFAP2C and PRDM1 with 389 common targets in hPGCLC (Figure S3B). In summary, GRN analysis highlighted how hiPSC and hEGCLC have a similar regulatory logic in term of key TFs and targets genes, suggesting MYC, ZIC5, HDAC2 and SP3 as key regulators and how these regulatory networks differ from that of hPGCLC, in which TFAP2C, PRDM1, KLF4 and REST emerged as master regulators.

### Longitudinal DNA methylation profiling to investigate the reversibility of epigenetic states

During their migratory phase, hPGCs initiate a wave of global DNA demethylation, including significant loss of methylation at imprinting control regions ^5–7^. To evaluate the extent to which our in vitro model recapitulates the genome-wide reprogramming events occurring in vivo, we profiled its DNA methylation levels (both 5-methylcytosine and 5-hydroxymethylcytosine) longitudinally (at least in triplicate for each cell type), by enzymatic methyl sequencing (EM-seq) ^86^. As evident from the PCA plot, samples corresponding to the same differentiation step cluster together, with the first principal component (PC1) explaining 36% of the variance mainly driven by hPGCLCs (Figure 4A). hiPSCs and hEGCLCs do not separate along the first principal component (PC1) but do so along the second one (PC2) that explains 19% of the variance (Figure 4A).

**Figure 4.**
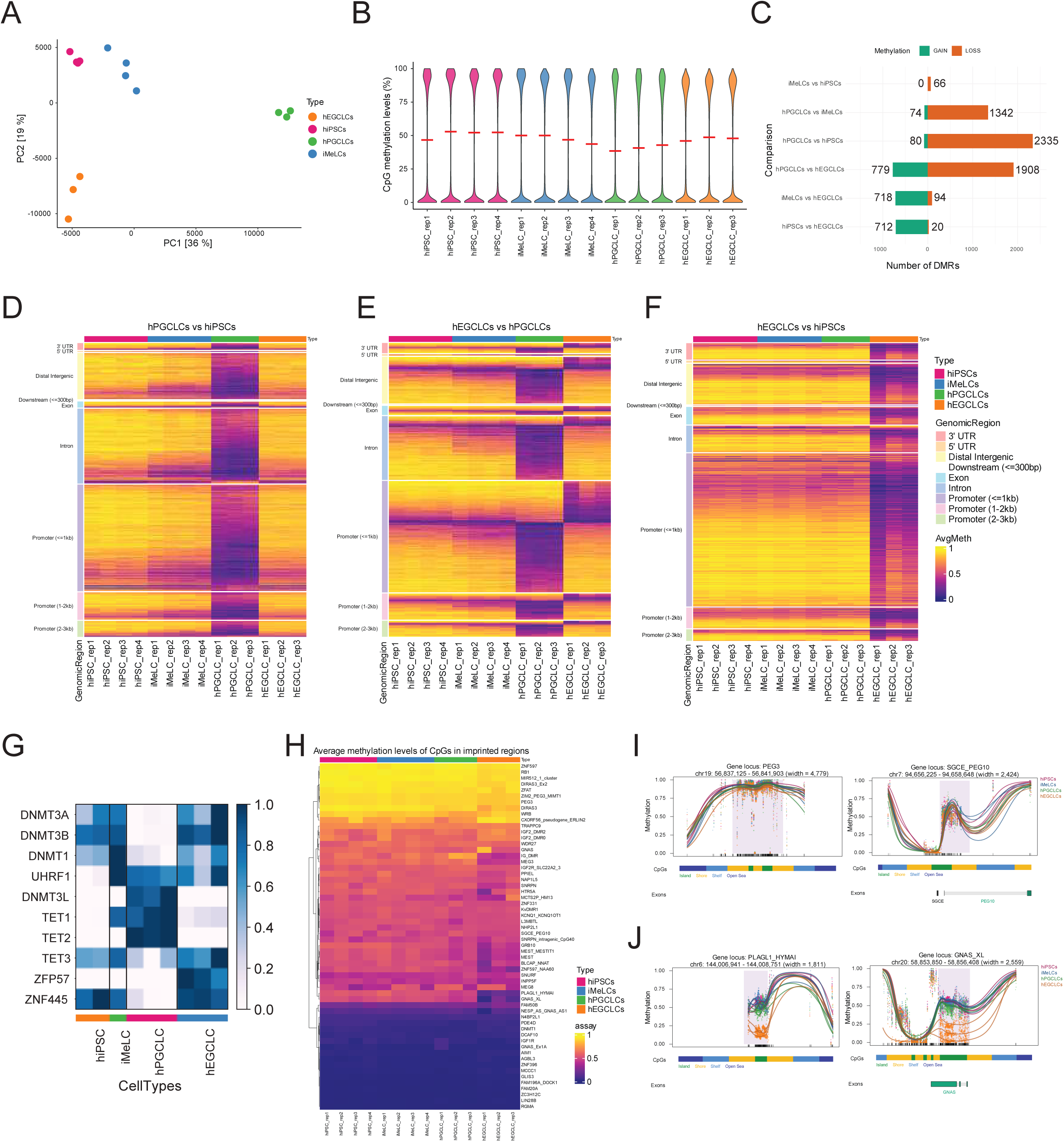
Longitudinal DNA methylation profiling to investigate the reversibility of epigenetic states **(A)** Principal component analysis (PCA) plot of CpG methylation levels of hiPSCs, iMeLCs, day 6 hPGCLCs and hEGCLCs (each cell type at least in triplicate). PC1 and PC2 explain 36% and 19% of the variability, respectively. **(B)** Violin plots showing the distribution of single CpG methylation levels (as percentage, Y-axis) at each measured region in each sample, X-axis. Methylation levels are defined as the number of methylated reads mapped at that CpG locus divided by the total number of reads mapped at that CpG locus. Red horizontal bars indicate the median methylation level. **(C)** Bar plots showing the number of differentially methylated regions (DMRs, X-axis) for each comparison between the four key cell types in pairs (Y-axis). Green bars indicate the number of DMRs that gain methylation in the first term of the comparison, orange bars the number of DMRs that lose methylation in the first term of the pairwise comparison. **(D), (E), (F)** Heatmaps of CpG average methylation level for each DMR (on the rows), for each of the samples analyzed (columns: hiPSCs and iMeLCs in quadruplicate, day6 hPGCLCs and hEGCLCs in triplicate). **(D)** hPGCLCs vs hiPSCs comparison; **(E)** hEGCLCs vs hPGCLCs comparison; **(F)** hiPSCs vs hEGCLCs comparison. Color gradient indicates the level of CpG methylation defined in (B). DMRs are clustered on the rows according to genomic region annotation depicted in the figure legend. **(G)** Heatmap of pseudo-bulk expression level in hiPSC, hPGCLC and hEGCLC of key genes involved in DNA (de)methylation and imprinting regulation. **(H)** Heatmaps of average methylation levels of CpGs belonging to imprinted genes (rows) for each of the samples analyzed (columns of the heatmap: hiPSCs and iMeLCs in quadruplicate, day6 hPGCLCs and hEGCLCs in triplicate). **(I)** and **(J)** DMR plots of four imprinted regions. Each dot represents the methylation level (Y-axis) of an individual CpG locus in a single sample, and its size is proportional to the coverage. The lines represent smoothed methylation levels for each sample (hiPSCs, iMeLCs, hPGCLCs, hEGCLCs). Gene exons and CpG annotations are shown below the plot if in the nearby.

Violin plots showing the percentage of global genomic DNA (gDNA) methylation for each sample suggests a gradual decrease in median methylation level along differentiation (hiPSC ◊ iMeLC ◊ day 6 hPGCLC), with a variation in median methylation levels between hiPSCs and hPGCLCs of about 20% (Figure 4B and Table S5),consistent with previous reports ^38,41^. Instead, compared to hPGCLCs, median gDNA methylation increases in hEGCLCs, reaching levels just slightly below those of hiPSCs, thus indicating that the approximately 20% global gDNA demethylation in hPGCLCs is largely reversible.

Next, we focused on the regions that, in pairwise comparisons between the cell types of our in vitro model, show significant DNA methylation differences (i.e. greater than 25%), therefore selected as differentially methylated regions (DMRs), as described in the Methods. Bar plots showing the total number of DMRs for each comparison (divided between gain and loss of methylation in one cell type compared to the other) confirm that hPGCLCs present lower methylation relative to all other cell types (Figure 4C). This is also evident by plotting CpG mean methylation levels for each DMR in heatmaps (Figure 4D, 4E and 4F) and violin plots (Figure S4A, S4B and S4C), for each comparison, grouping DMRs according to the corresponding genomic region annotation (i.e. promoter, 5’ and 3’ UTRs, exon, intron, downstream [of gene end] and intergenic regions). Interestingly, methylation trends are similar across all these genomic regions for all the comparisons (Figure S4A, S4B and S4C and Table S5). Considering the DMRs from hiPSC-hPGCLC comparison (Figure 4D), hEGCLCs have CpG methylation levels similar to hiPSCs, confirming the similarity between hiPSCs and hEGCLCs when compared to hPGCLCs and the reversibility of the hPGCLC demethylation wave ^41^. Examples of regions (i.e. promoter region of DNMT3L ^7,87^, MCF2L and CSF3R genes ^81^) showing a significant loss of methylation in hPGCLCs compared to hiPSCs and hEGCLCs are depicted in Figure S4D. Functional enrichment analysis performed on genes associated to regions significantly differentially methylated in hPGCLCs compared to hiPSCs or hEGCLCs highlights terms mainly related to regulation of cell shape, histone deacetylation and regulation of hippo signalling (Figure S4E and Table S5).

To distinguish between 5-methylcytosine (5mC) and 5-hydroxymethylcytosine (5hmC), we quantified their level in our longitudinal dataset by LC-MS/MS (see Methods). This analysis confirmed a slight decrease of 5mC levels in hPGCLCs compared to hiPSCs and hEGCLCs (Figure S4F top panel) and highlighted an increase of 5hmC levels in hPGCLCs compared to the pluripotent cell types, that for both cytosine modifications exhibit similar levels (Figure S4F bottom panel). This is in line with the initiation of methylome resetting in nascent hPGCLCs via oxidation of 5mC to 5hmC by TET enzymes at certain loci ^5,88,89^.

We also examined the expression of genes involved in the regulation of DNA methylation in our scRNAseq dataset. DNA methyltransferases (DNMT1 and DNMT3A/B) expression decreases in hPGCLCs compared to hiPSC but recovers in hEGCLCs. In contrast, the expression of TET1 and TET2 demethylase and of DNMT3L is low in hiPSCs and hEGCLCs and increases in hPGCLCs (Figure 4G). However, we were surprised to find that the expression of the DNMT1 cofactor UHRF1 increases as hiPSCs differentiate to hPGCLCs but does not decrease in hEGCLCs (Figure 4G).

Finally, we explored specific DMRs, of which we identified 712 hypomethylated and 20 hypermethylated regions in hEGCLCs compared to hiPSCs (Figure 4C, 4F). Functional enrichment analysis performed on hypomethylated regions highlighted terms related to cell adhesion and regulation of small GTPase mediated signal transduction (Figure S4G). Intriguingly, the promoter of PIWIL2 is amongst these hypomethylated regions, in keeping with its increased expression in hEGCLCs compared with hiPSCs (Figure S4H). These hypomethylated regions emerge as pluripotent stem cells passage through a germ cell state and therefore may reflect an epigenetic memory of this cell fate transition.

With this in mind, we next focussed on regions subjected to genomic imprinting. Loss of DNA methylation at imprinting control regions has been observed in many mouse EGC lines ^20,90,91^, although EGC lines derived from early PGCs can emerge with imprints intact ^21^. Global methylation levels at the different human imprinted regions are similar in hiPSCs, iMeLCs, hPGCLCs and hEGCLCs (Figure 4H and Table S6). This pattern is exemplified by PEG3 ^92,93^ and PEG10-SGCE ^94,95^, the two imprinted regions with the highest coverage in terms of number of CpG detected in our dataset (Figure 4I). As such, hPGCLCs largely recapitulate early hPGC development (around post-conceptional week 2-3), when imprinted regions have not yet undergone DNA demethylation ^11,40,43,96–98^.

Interestingly, however, we noted that hPGCLCs downregulate ZFP57 and ZNF445 (Figure 4G), two regulators critical for imprinting maintenance ^99^, and that selected imprinted regions, including PLAG1-HYMAI ^100^, GNAS ^101–103^ (Figure 4J) and MEST ^104^, did exhibit lower levels of methylation in hEGCLCs when compared with hiPSCs and hPGCLCs. These imprinted regions overlap with those that have recently been reported to be demethylated during the early stages of hPGCLC culture ^105^. It is therefore possible that these loci are more susceptible to DNA demethylation, or that factors used to culture hPGCLCs in vitro play a role in triggering their demethylation. We also note that imprint instability in culture is a well-described phenomenon during pluripotent stem cell culture ^106,107^. The extent to which loss of DNA methylation at imprinted regions during in vitro culture of PGCs/PGCLCs accurately reflects the in vivo process of imprint erasure and whether this can be manipulated by altering the culture environment are important areas for future study. Overall, these data confirm that the initial demethylation associated with hPGCLC induction is largely reversible, but that a small number of regions of potential relevance to germ cell identity do not regain DNA methylation.

### Multi-omic gene regulatory network analysis identifies key regulators of the pluripotency-germline transitions

Leveraging the multi-omics nature of our longitudinal dataset, we integrated DNA methylation and transcriptomic data to further dissect the master regulators governing the pluripotency-germline transitions. Focusing on genes exhibiting an inverse correlation between methylation levels at their regulatory regions and their expression we specifically examined those that were up- or down-regulated in hPGCLCs compared to pluripotent cells (both hiPSC and hEGCLC) (Figure 2D, 5A).

Master regulator analysis (MRA) on these differentially methylated and expressed genes (dMdE) identified 136 key TFs (Figure 5B, S5B). Interestingly, while only 7 of these TFs emerged as key regulators across all 4 cell types, we observed that subsets of the TFs exhibited cell-type specificity, with 52 TFs, including members of the GATA, SOX ^68^, and ZNF families ^5^, specific for hPGCLCs, 10 TFs specific for hiPSC (IRX3, NFATC4, ATOH8, PRDM11, PKNOX2, NR5A2, TBX3, MEF2B, GBX2, HDAC2), and 4 specific for hEGCLC (SOX18, ZNF394, RXRG, MYC). This observation confirms our previous identification, from the transcriptomic-specific analysis, of MYC and HDAC2 as top regulators of pluripotency networks. However, despite the similarities between hiPSC and hEGCLC GRNs, key regulatory differences exist, particularly in genes that display dynamic expression and methylation profiles during the reprogramming process (Figure S5A).

**Figure 5.**
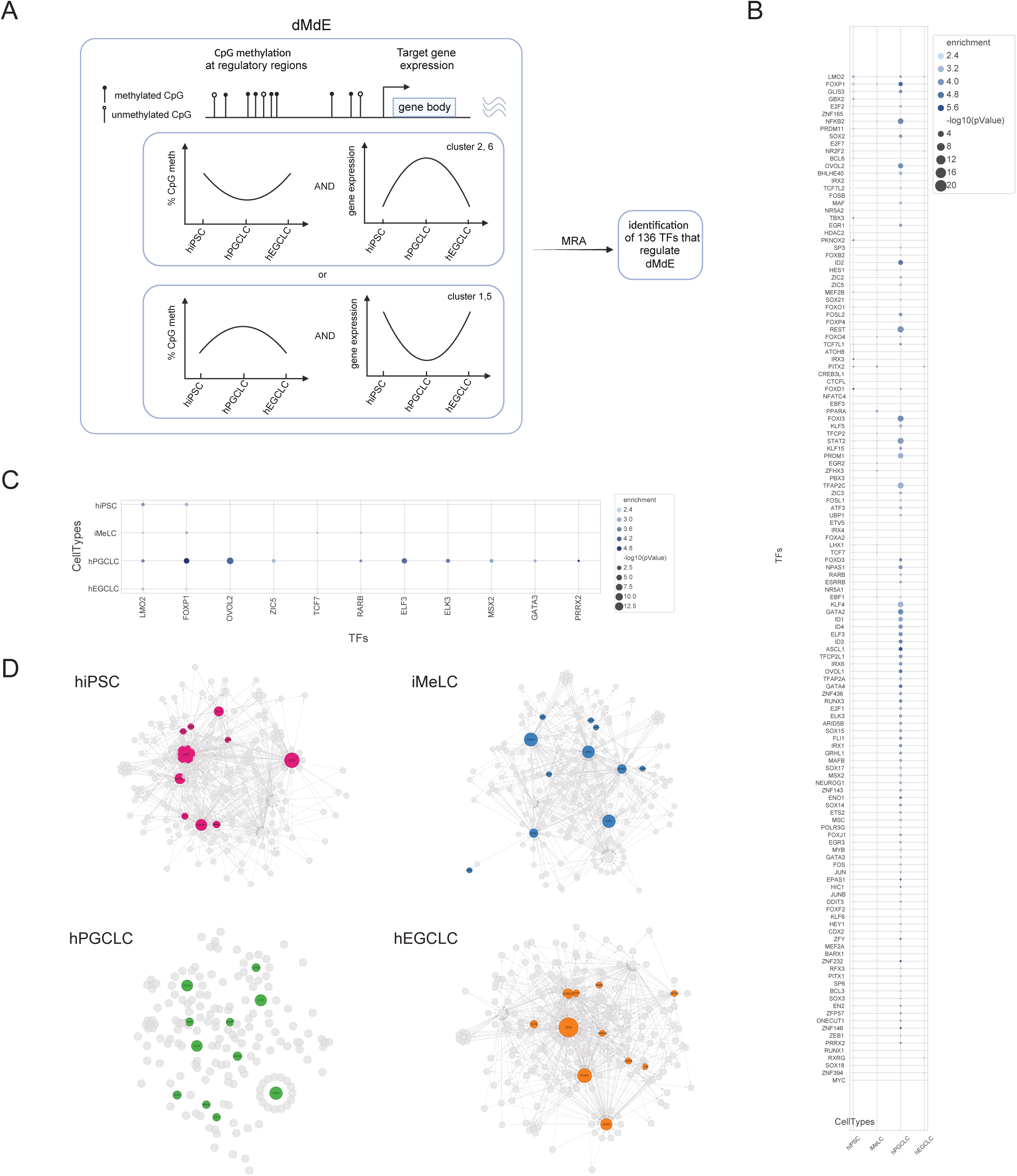
Multi-omic gene regulatory network analysis identify key regulators of the pluripotency-germline transitions **(A)** Schematics of the selection strategy for dMdE genes by integration of DEGs and DMRs. MRA was then performed to identify TFs responsible for the regulation of these genes. **(B)** Dot plot of TFs (Y-axis) significantly regulating dMdE genes in each cell type (X-axis), (enrichment>2; -log10(p-value)>1.3, p-value< 0.05). **(C)** Dot plot of dMdE TFs (X-axis) significantly regulating dMdE genes in each cell type (Y-axis), (enrichment>2, p-value< 0.05). Dot color represents enrichment and dot size represents -log10(p-value). **(D)** GRNs visualization (one for each of the four cell types), highlighting the 11 TFs (colored) that are differentially expressed and differentially methylated themselves and that regulate dMdE genes; their targets and regulators are in grey.

Furthermore, our analysis revealed that among the differentially methylated and expressed genes (dMdE), 11 were themselves transcription factors that significantly regulated other dMdE genes (Figure 5C, 5D, S5C), and 17 were methylation-sensitive TFs, namely TFs for which the methylation state of their recognition sequence can directly modulate their binding affinity ^108–111^ (Figure S5D and Table S7).

These findings highlight the intricate interplay between DNA methylation and transcriptional regulation during germline differentiation and reprogramming, and underscore the importance of considering both epigenetic and transcriptomic dynamics to study these processes.

### Escapee regions are hypermethylated compared to non-escapee regions and resistant to reprogramming

Previous studies reported “escapee” regions, that exhibit resistance to DNA demethylation in both mouse ^112^ and human PGCs ^5^ and proposed these as candidates which might mediate epigenetic inheritance ^5^. Our protocol presents a unique opportunity to study how these epialleles behave during germline induction and subsequent reprogramming to hEGCLCs. Among the 3,680,616 CpG loci analysed, 9113 fall within the escapee regions identified by Tang et al. ^5^. A PCA plot of CpG methylation levels at escapee regions shows how hiPSCs and hPGCLCs separate along the first PC (explaining 20% of the variance), however conversion to hEGCLCs is not accompanied by a reversion in methylation, as hEGCLCs cluster together with hPGCLCs along PC1. hPGCLCs and hEGCLCs separate only along the PC2 (that explain 16% of the variance), as do hiPSCs (Figure 6A). This contrasts markedly with the PCA plot of non-escapee regions (Figure S6A). The non-escapee pattern is almost identical to the combined PCA plot (Figure 4A), as expected, since the majority of the CpG loci belong to non-escapee regions.

**Figure 6.**
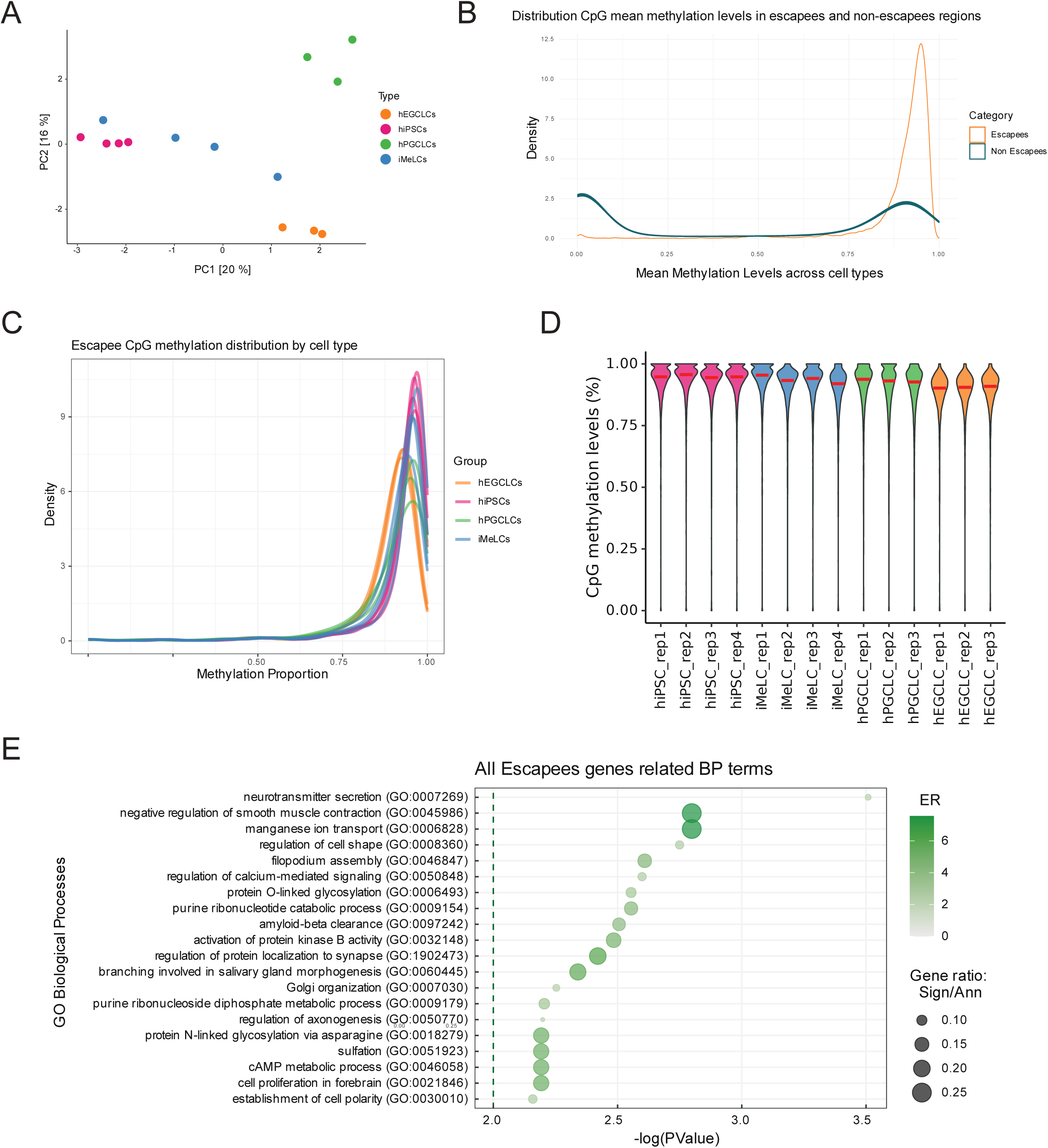
Escapee regions are hypermethylated compared to non-escapee regions and resistant to reprogramming **(A)** Principal component analysis (PCA) plot of CpG methylation levels of escapee regions of hiPSCs, iMeLCs, day 6 hPGCLCs and hEGCLCs. PC1 and PC2 explain 20% and 16% of the variability, respectively. **(B)** Density plot of methylation level distributions of the 9113 escapee CpGs and of 500 sets of randomly selected CpGs within the non-escapee regions (cell types collapsed by average). **(C)** Density plot of the methylation level distribution of the 9113 escapee CpGs, colored by cell type. **(D)** Violin plots of CpG methylation levels of escapee regions (%, Y-axis) for each sample (X-axis). Methylation levels are defined as the number of methylated reads mapped at that CpG locus divided by the total number of reads mapped at that CpG locus. Red horizontal bars indicate the median methylation level. **(E)** Dot plot of enriched biological process (BP) gene ontology (GO) terms for the genes annotated to the escapee regions. Y-axis: GO term; X-axis: -log(p-value). Color gradient indicates the enrichment ratio (ER)>2, dot dimension indicates gene ratio. The dotted vertical line indicates the p-value threshold at 0.01.

We then compared the distribution of CpG mean methylation levels in escapees vs non-escapee regions (Figure 6B), noting how non-escapee regions follow a bimodal distribution (Figure 6B and S6B), as expected, whereas escapee regions follow a unimodal distribution, shifted towards higher methylation levels (Figure 6B and 6C). This is consistent with the tendency of these regions to be hypermethylated and to escape demethylation ^5,113^. When looking at escapee regions CpG methylation distribution by cell type (Figure 6C), we noticed how, even if all the four cell types of our in vitro model present a unimodal distribution skewed towards higher values, globally methylation levels are lower in hPGCLCs compared to hiPSCs and even lower in hEGCLCs (Figure 6D, S6C). This result points to a peculiar behaviour of these regions during the conversion of hPGCLCs to hEGCLCs. While the majority of genomic loci re-accumulate DNA methylation to levels similar to hiPSCs, the escapee regions remain relatively demethylated. Thus, these regions are not simply characterised by an inherent propensity to maintain high levels of DNA methylation. Rather they exhibit a context dependent response to different forms of epigenetic reprogramming - remaining relatively hypermethylated during epigenetic reprogramming in late PGCs, while remaining relatively hypomethylated during the generalised reacquisition of DNA methylation during EGCLC derivation.

Functional enrichment analysis of the genes associated with escapees-regions included in our analysis shows an enrichment for terms related to neurodevelopment, such as regulation of protein localization to synapse and of axogenesis (Figure 6E), in keeping with the findings by ^5^. As this original study specifically raised the possibility of intergenerational epigenetic inheritance at these regions, our finding that escapee regions might be specifically protected from remethylation during PGCLC-to-EGCLC conversion, may suggest context-dependent regulation of DNA methylation at these loci and establish our culture system as a new paradigm to investigate the targeted alteration and inheritance of clinically relevant epialleles.

## Discussion

Here we report the first fully defined system to track the pluripotent state transitions between PSCs and hPGCLCs. We demonstrate that hPGCLCs can efficiently convert back to the PSC state from which they are derived, both at the functional and molecular level, using multiple hPSC lines. Our multi-omic tracking of these transitions provides a high-resolution map of the transcriptional and epigenomic transitions upon entry to and exit from the human germline.

In mice since the first reports of EG cell derivation there has been significant progress in identifying factors that either promote PGC proliferation and/or their conversion to EG cells (reviewed in ^2^). These advances led to the development of a fully defined system to culture mouse PGCs and track their conversion to EG cells ^22^. In our experience, mouse PGCLCs readily convert to form EGCLCs in the same conditions (LPS-R and HGL, unpublished observations) and modifications of the PGC culture system have enabled long term expansion cultures of mouse PGCLCs ^114^. Despite these advances in the mouse, bona fide human EG cells derived from hPGCs have not yet been reported. While recent studies have highlighted the differences between mouse and human PGC development, there are many similarities between PGCs from mice and humans, both in vivo and in vitro ^115^. Importantly, the same factors used to trigger mPGCLC induction are also effective in human PGCLC ^3^. Our findings establish that the same factors that support the derivation of mouse EG cells also trigger the efficient transition of hPGCLCs to hEGCLCs. Moreover, the similar culture conditions for PGCLC-to-EGCLC between mice and humans suggest that the mechanisms of conversion might be evolutionarily conserved between mammals - an exciting avenue for future investigation.

There has been a previous report of hEGCLC derivation ^41^. In this study, the authors initiate hPGCLC cultures on a feeder cell layer and then transition the cells to Matrigel, in a feeder-cell conditioned medium. This system enables an impressive degree of hPGCLC expansion in long term cultures over hundreds of days. Subsequent removal of conditioned medium and culture with FGF2 and SCF leads to the formation of pluripotent stem cell colonies which can be picked and expanded as hEGCLC lines. Intriguingly, a recent report noted that dedifferentiation can occur in long term hPGCLC cultures maintained on feeder cells, albeit using different culture medium ^105^. Thus, it seems possible that feeder cells produce a range of factors which variably either promote or inhibit the emergence of hEGCLCs.

Rather than obtaining hEGCLCs as they emerge from long term PGCLC cultures, we have established a dedicated hEGCLC derivation protocol. Notably our hEGCLC derivation medium does not support long term expansion of hPGCLCs. This is in keeping with the idea that unknown factors produced by feeder cells and present in conditioned medium are key to this long-term expansion. The system reported here may be one route to screen for such factors as, unlike other systems, cultures are initiated and maintained without feeders or feeder cell-conditioned medium. This will facilitate future studies to identify the pathways that enable hPGCLCs to transition to a PSC state. Future work should address whether hPGCLCs can also give rise directly to naive PSCs, helping to deepen our understanding of pluripotent states in humans - which at the current time is less well understood than in the mouse.

Our defined culture system facilitated multi-omics longitudinal profiling which in turn allowed us to capture germline transcriptional program emergence and dissolution. While there is substantial transcriptomic similarity between hiPSCs and hEGCLCs, we identified a small subset of genes that are upregulated in hPGCLCs but do not return to their previous hiPSC-levels in hEGCLCs. In keeping with previous findings ^41^, this included the germ cell specific gene PIWIL2. Our data demonstrates that this transcriptional memory of transition through a germline state coincides with failure to re-establish DNA methylation at PIWIL2’s promoter, suggesting that the gene expression differences may be driven by incomplete epigenetic resetting. It is possible that for some germline genes that are regulated by DNA methylation, targeted remethylation can only occur during a discrete time window - such as the accumulation of DNA methylation at germline specific genes that occurs upon implantation in the mouse ^116^. In keeping with a germline specific effect, we also observed non-reversible demethylation of a small number of imprinted regions. Intriguingly, we observed a similar phenomenon at previously reported ‘escapee regions’ - suggesting that this group of genes may exhibit more complex, context-dependent epigenetic regulation, rather than simply being resistant to DNA demethylation in PGCs. More broadly, our culture system may enable future studies to investigate the inheritance and targeted alteration of clinically relevant epialleles during transition through a germline state.

Our detailed multi-omic analysis of changes in gene expression and epigenetic state enabled us to describe the gene regulatory logic of the hPSC-to-hPGCLC-to-hEGCLC transition and identify key transcription factors that control these changes in cell state. This knowledge may have significant applications in both the stem cell and cancer fields. For instance, our study represents an important step towards the derivation of hEGC from in vivo hPGC. Our defined protocol will allow us to attempt hEGC derivation in an iterative and targeted way. For instance, by altering media composition to more specifically regulate the expression of TFs and pathways that drive the hPGCLC to hEGCLC transition. Access to early stage hPGCs is likely to remain a major challenge, although such samples do rarely become available to researchers ^54^. The derivation of human EGC would be a major breakthrough in the field, establishing that the ability to derive the full repertoire of embryo-derived PSCs is not a rodent-specific phenomenon. This in turn will help us better understand the different pluripotent states in humans - including how well they map to the naive, formative, primed and latent states that are increasingly well defined in mice ^24,117– 120^.

An understanding of how human PGCs transition to a PSC state is also of clear relevance to our understanding of the pathogenesis of germ cell tumours (GCTs) in humans - many of which are derived from hPGCs and have pluripotent and stem cell characteristics. In particular, sacrococcygeal teratomas represent the commonest neonatal tumours in humans ^121^ and germ cell tumours amongst the most common childhood cancers ^122^. Variants in SPRY4, GAB2, and KITLG are associated with an increased risk of GCTs and it is noteworthy that each of the associated signalling pathways are targeted by factors in our hEGCLC derivation medium ^123^. A role for environmental factors, including endocrine disrupting chemicals, has also been proposed^124^. Our culture system therefore represents a new method to screen potential genetic, signalling, and chemical triggers that might trigger GCT formation in humans. In addition, it seems possible that the TFs we have identified as drivers of the hPGCLC-hEGCLC transition might be relevant to GCT pathogenesis, a proposal that merits future investigation.

As a whole, this multi-omics longitudinal profile will be a key resource for the scientific community to leverage on and exploit in several applications domains, from deepening the understanding of pluripotent states in humans, to the study of the genetic and environmental factors involved in GCT pathogenesis and to the investigation of the epigenetic dynamics occurring during the germline cycle.

## Material and Methods

### Human iPSC lines culture

Healthy controls-derived hPSC lines were cultured under feeder-free conditions on matrigel (Corning, #354277) or Geltrex (Life Technologies, A1413302) coated plates at 37°C, 5 % CO2 and 5 % O2. The hiPSC cell line BTAG (585b1-868) was a kind gift from Prof Mitinori Saitou (Institute for the Advanced Study of Human Biology, Kyoto, Japan). The SOX2-tdTomato cell line was a kind gift from Prof. Timo Otonkoski (University of Helsinki, Norway). Before plating cells, dishes were coated with matrigel solution (Corning, #354277, according to the manufacturer’s instructions). hiPSCs were maintained in TeSR/E8 medium (Stemcell Technologies, #05990) or StemFit medium (Amsbio #SFB-504-CT), supplemented with 100 U/mL penicillin and 100 ug/mL streptomycin (ThermoFisher, #15140122) with daily media changes and splitted 1:8 to 1:10 with ReLeSR (Stemcell technologies, #05872) when confluency reached around 70%. When single cell dissociation was needed, Accutase (Sigma-Aldrich, #A6964) was used instead of ReLeSR, and 10uM ROCK inhibitor Y-27632 (Tocris, #1254) was added to the medium to enhance cell survival in the first 24 hours. All experimental activities involving hPSC were approved by the ethical committee of the University of Milan. All hPSC lines were reprogrammed by at least 15 passages, verified to be mycoplasma free by routine PCR testing and their identity was confirmed by short tandem repeat profiling. Details about the hPSC lines can be found in the table below:

| In house name | Cell line ID | Source | Type | Sex | Origin | Profiling done in this study |
| --- | --- | --- | --- | --- | --- | --- |
| **BTAG** | 585B1-868 | Sasaki et al., 2015 | hiPSC | M | Peripheral blood, healthy donor | Live imaging, IF*, tri-germ**, bulkRNAseq, LC-MS/MS |
| **SOX2-tdTomato** | UHi001-A | University of Helsinki (UH), Balboa et al., 2017 | hiPSC | M | Fibroblast, healthy donor | Live imaging |
| **CTL08A** | WTSli018-B-1 | Wellcome Sanger Institute (WTSI) | hiPSC | M | Fibroblast, healthy donor | IF*, tri-germ**, scRNAseq, EMseq |
| **H9** | WAe009-A | WiCell Research Institute (WA) | hESC | F | Blastula, with ICM and trophoblast, no disease diagnosed | IF* |
| **CTL04E** | UMILi024-A | University of Milan (UMIL) | hiPSC | F | Fibroblast, healthy donor | IF*, tri-germ** |

\*IF = Immunofluorescence staining for pluripotency markers
\*\*tri-germ = tri-germ layer differentiation

### Induction of iMeLCs and hPGCLCs

hPGCLCs were induced from hiPSCs via iMeLCs as described in ^1,2^, with minor modifications. Briefly, iMeLCs were induced by plating 0.5-1.0 x 10^5^ hiPSCs maintained in StemFit or TeSR/E8 onto a well of a human plasma fibronectin (Sigma-Aldrich, #F0895-5MG)-coated 12-well plate in GK15 medium [GMEM (Thermo Scientific, #11710035) with 15% of knockout serum replacement, KSR (ThermoFisher Scientific, #10828028), 0.1 mM Non-essential amino acids, NEAA (Sigma, #M7145), 2 mM GlutaMax (Gibco, #35050061), 1 mM sodium pyruvate (Thermo Scientific, #11360070), 0.1 mM cell culture grade 2-mercaptoethanol solution (Gibco, #31350010) and penicillin 100 U/mL and streptomycin 100 μg/mL (ThermoFisher, #15140122)] containing 50 ng/ml of Activin A (Peprotech, #120-14P), 3 µM of CHIR99021 (Sigma-Aldrich, #SML1046-5MG), 20 nM PD173074 (Stemgent, #04-0008, only for CTL08A and CTL04E cell line), and 10 µM of a ROCK inhibitor Y-27632 (Tocris, #1254). hPGCLCs were induced by plating 3000-5000 iMeLCs dissociated in single-cell into a well of a ultra-low attachment, V-bottom 96 well plates (SystemBio, #MS-9096VZ) in GK15 supplemented with 10 ng/ml of LIF (PeproTech, #300-05), 200 ng/ml of BMP2 or BMP4 (qkine, #Qk007-0100 and Qk038-0100, respectively), 100 ng/ml of SCF (PeproTech, #300-07), 50 ng/ml of EGF (Peprotech, #AF-100-15), and 10 µM of ROCK inhibitor.

### Fluorescence-activated cell sorting

At day 4 or day 6 up to 24 floating aggregates were collected per 1.5-ml tube, washed twice in PBS and dissociated with 800ul/tube 0.25% Trypsin-EDTA in PBS (Thermo Fisher Scientific, #15090046) for 15-20 min at 37°C, with periodical intermittent pipetting. The cell suspension was quenched with PBS containing 10% FBS (Thermo Scientific, #26140079) and 0.1% bovine serum albumin (BSA), followed by pipetting to generate a single-cell suspension. The cell suspension was then subjected to centrifugation for 4 min at 200 rcf and the pellet resuspended in 1ml GK15 and filtered by using 70-micron Flowmi cell strainers for 1000 ul tips (Merck, #BAH136800070). When not using the BTAG reporter line ^1^, cells were stained with FITC-conjugated anti-human CD326 (EpCAM) antibody (BioLegend, #324204) and PE-conjugated anti-human/mouse CD49f antibody (Miltenyi Biotec, #130-119-767) for 15 minutes at room temperature (RT). hPGCLCs were sorted and isolated with a cell sorter (Beckman Coulter, MoFlo Astrios Cell Sorter), and collected in GK15 medium plus 10 µM Y-27632.

### hEGCLCs derivation from sorted hPGCLCs

To reprogram hPGCLCs into hEGCLCs, sorted day 4 or day 6 hPGCLCs were plated directly into 96 multi-well flat bottom (10 cells/well), pre-coated with Geltrex (Thermo Scientific, #A1413201), according to the manufacturer’s instructions, and containing 100 ul/well of hEGCLC induction medium [i.e. N2B27 medium ^3^ containing 10 ng/ml human leukemia inhibitory factor (hLIF, PeproTech, #300-05), 3 µM of the GSK3 inhibitor CHIR99021 (CH, Sigma-Aldrich, #SML1046-5MG), 10 µM Forskolin (FK, Sigma-Aldrich, #F3917-25MG), 25ng/ml basic fibroblast growth factor (bFGF, Preprotech, #100-18B), 100 ng/ml human stem cell factor (hSCF, PeproTech, #300-07) and 2 µM retinoic acid (RA, Sigma-Aldrich, #R2625)] and antibiotics (penicillin 100 U/mL and streptomycin 100 μg/mL, ThermoFisher, #15140122). They were grown at 37 °C, 5 % CO2 and 5 % O2. After 48 hours 100ul of StemFit (Amsbio, #SFB-504-CT) were added to each well. On day 3, 100 ul of medium were removed and added 100ul of StemFit. On day 4, 150 ul of medium were removed and 100ul of StemFit added. On day 5, all medium was removed and 100ul of StemFit added. From day 6 onwards, total media change was performed every other day. In these defined and feeder-free culture conditions 10-15% of hPGCLCs undergo conversion to form hEGCLC colonies over a period of two weeks (conversion efficiency was calculated as total number of hEGCLC colonies obtained divided by total number of hPGCLC seeded [ie. 10 x total number of well seeded, since we plated 10 hPGCLC/well]). These hEGCLC colonies were then picked, expanded and maintained in conventional human pluripotent stem cell conditions.

### Immunofluorescence (IF) analysis

hEGCLCs were seeded in 24 multi-wells coated with Matrigel and grown in StemFIt medium. When around 60% confluency was reached, colonies were washed three times with PBS and fixed with 4% paraformaldehyde/PBS solution (PFA, SantaCruz, #sc-261692) for 15 minutes at RT. Then cells were washed three times with PBS and permeabilized with 0.5% TRITON X-100 in PBS for 15 minutes at RT. Blocking was then performed using 0.3% TRITON X-100 + 5% Donkey Serum (DS, (Jackson ImmunoResearch, #017-000-121) in PBS 1X for 60 min at RT. Incubation with primary antibodies was performed in 0.2% TRITON X-100 + 2% DS in PBS at 4°C overnight. The following primary antibodies were used: OCT4 (Rabbit, 1:500, Abcam #ab181557), SOX2 (Goat, 1:200, R&D System #AF2018), NANOG (Goat, 1:400, Everest Biotech #EB06860). The day after, three washes in PBS were performed, followed by RT 120 minutes incubation with secondary antibodies conjugated with AlexaFluor PLUS 555 or 647 (Thermo Fisher Scientific) diluted 1:400 in 0.2% TRITON X-100 + 2% DS in PBS. Nuclei were stained with Hoechst 33342 (Cell Signaling Technology) 0.1 μg/ml in PBS 1X for 10 min at RT. After this incubation three washing steps with PBS were performed. Cells were left in PBS + 0.02% Sodium azide. Images at 10X and 20X magnification were acquired using an Axio Vert.A1 FL-Inverted LED fluorescence microscope (Zeiss) equipped with Axiocam 712 mono CMOS camera and then processed with ImageJ Fiji image analysis software.

### Evaluation of pluripotency markers expression by flow-cytometry

hEGCLC pluripotency markers expression was evaluated also by flow-cytometry using BD Stemflow™ Human Pluripotent Stem Cell Transcription Factor Analysis Kit (#560589), following manufacturer instructions. Stained single cell suspensions were analyzed using a Beckman/Coulter CytoFlex LX Analyzer. Data were analyzed using FCS-Express 7 software.

### Three germ layer differentiation and immunofluorescence

hEGCLCs were collected from one 70–80% confluent 6 cm dish and plated in a low attachment 35 mm dish (S-Bio, # MS-90350Z) in 1 ml of exhausted medium (StemFit; Amsbio, #SFB-504-CT) and 1 ml of fresh StemFit + 10 mM Y-27632. After 2 days media was changed with 1 ml StemFit + 1 ml complete KSR EB Medium (1x DMEM/F12, 100 U/mL penicillin and 100 ug/mL streptomycin, 2mM GlutaMAX, 20% KSR, 0.1 mM Non-Essential Amino Acids). After 2 days the media was changed with 2 ml complete KSR EB Medium. After 2 days, up to three embryoid bodies (EBs) were seeded in each well of a geltrex-coated 24 multi-well in complete KSR EB Medium, allowing them to attach and grow. Media was changed every other day until day 21 when they were washed three times with PBS and fixed adding 300ul of 4% paraformaldehyde/PBS solution (SantaCruz, #sc-261692) for 15 minutes at RT. Then EBs were washed three times with PBS and permeabilized with 0.5% TRITON X-100 in PBS for 15 minutes at RT. Blocking was then performed using 0.3% TRITON X-100 + 5% Donkey Serum (DS, (Jackson ImmunoResearch, #017-000-121) in PBS 1X for 60 min at RT. Incubation with primary antibodies was performed in 0.2% TRITON X-100 + 2% DS in PBS o/n at 4°C. The following primary antibodies were used: FOXA2 (mouse, 1:400, Abcam #ab60721), SOX17 (rabbit, 1:400, Millipore #09-038-I), PAX6 (rabbit, 1:400, Biolegend #901301), MAP2 (guinea pig, 1:400, Synaptic Systems #188 004), TUBB3 (mouse, 1:400, BioLegend #801202), α-SMA (mouse, 1:1000, Sigma-Aldrich #A5228). The day after, three washes in PBS were performed, followed by RT 120 minutes incubation with secondary antibodies conjugated with AlexaFluor PLUS 555 or 647 (Thermo Fisher Scientific) and Goat anti Guinea Pig Secondary Antibody, Alexa Fluor 647 (Thermo Scientific, A-21450) diluted 1:400 in 0.2% TRITON X-100 + 2% DS in PBS. Nuclei were stained with Hoechst 33342 (Cell Signaling Technology) 0.1 μg/ml in PBS 1X for 10 min at RT. After this incubation three washing steps with PBS buffer were performed. Cells were left in PBS + 0.02% Sodium azide. Images at 10X and 20X magnification were acquired using an Axio Vert.A1 FL-Inverted LED fluorescence microscope (Zeiss) equipped with Axiocam 712 mono CMOS camera and then processed with ImageJ Fiji image analysis software.

### Live imaging

Time-lapse imaging was performed on an Olympus IX83 microscope equipped with a Hamamatsu ORCA-Flash 4.0 camera and environmental chamber kept at 37°C with a 5% CO_2_ supply. Pictures were collected every 3 hrs for 120 hrs using a UCPlanFN N 40x/0.60 objective lens. Ten hPGCLCs were placed per well of a 96-well plate (655090, Greiner Bio-One, previously coated with GelTrex) at the start of the hPGCLC-hEGCLC transition protocol and placed for time-lapse at different starting time points.

### RNA extraction and bulk RNA sequencing library preparation

Frozen pellets were submitted to GENEWIZ for RNA isolation, library preparation with poly(A) selection and 150bp paired end sequencing on Illumina.

### Single-cell dissociation and single-cell RNA sequencing library preparation

hiPSCs, iMeLCs, hEGCLCs were detached and dissociated into single-cell using Accutase (Sigma-Aldrich, #A6964). 96 day 6 hPGCLCs aggregates were collected and dissociated into single-cells as described in Fluorescence-activated cell sorting section. Single cells were resuspended in chilled 0.04% Bovine Serum Albumin (BSA, Merck, A9418-50G) in HBSS (ThermoFisher Scientific, #14025092), filtered by using 70-micron Flowmi cell strainers for 1000 ul tips (Merck, #BAH136800070) and counted with a Countess 3FL automatic cell counter (Thermo Fisher, #AMQUAF2000) using Trypan blue (Gibco, #15250062). Up to 1.5 million cells were taken from the cell suspension and processed according to 10X Fixation of Cells & Nuclei for Chromium Fixed RNA Profiling protocol (CG000478, Rev A). Briefly, filtered cells were centrifuged for 10 min at 300 rcf at 4°C and the pellet resuspended in 1 ml Fixation Buffer and incubated at 4°C. After 20 hours, fixed cells were centrifuged at 850 rcf for 10 minutes at RT and pellet resuspended in 1 ml chilled Quenching Buffer. After counting dead cells, 100 ul of pre-warmed Enhancer and 275 ul of 50% glycerol were added to fixed samples in Quenching Buffer. Samples were stored at -80°C. scRNAseq libraries were prepared following Chromium Fixed RNA Profiling for Multiplexed Samples 10X protocol (CG000527-RevC). Briefly, samples were thawed, centrifuged at 850 rcf for 5 minutes at RT and pellet resuspended in 1 ml 0.5X PBS + 0.02% BSA. Cell concentration of the fixed samples was then determined using a Countess 3FL automatic cell counter.

A total of 7000 cells per sample were processed, pooling together 16 individually barcoded samples, after the hybridization step. A total of 8 cycles of PCR was used for library amplification. Libraries were sequenced on a NovaSeq6000 platform (Illumina). Using proper cluster density, a coverage of at least 25,000 reads/cell was obtained. Cell Ranger version 7.0.0 (10X Genomics) was used to generate FASTQ files.

### DNA extraction and Enzymatic Methyl-seq library preparation and sequencing

Genomic DNA was extracted using DNeasy Blood & Tissue Kit (Qiagen, #69504), and libraries were synthesized from 200 ng purified DNA using NEBNext Enzymatic Methyl-seq (EM-seq) Library Preparation Kit (NEBNext, # 101977). Unmethylated Lambda phage DNA was spiked-in to evaluate the efficiency of bisulfite conversion. To generate target-enriched DNA libraries, Twist Targeted Methylation Sequencing Kit (TWIST Bioscience, #104181) was used, coupled with Twist Human Methylome Panel (TWIST Bioscience, #105521), according to the manufacturer’s instructions (DOC-001224 REV 4.0). Libraries were pooled in equimolar quantities and the pool of libraries was amplified by qPCR. Quantification and validation of enriched libraries was performed by Qubit dsDNA High Sensitivity Quantitation Assay (Thermo Fisher Scientific, #Q33230) and Agilent Bioanalyzer High Sensitivity DNA kit (Agilent Technologies, #5067-4626). Libraries were sequenced on a NovaSeq6000 platform (Illumina) with at least 160X coverage.

### Quantification of 5-hydroxymethylcytosine and 5-methylcytosine by LC-MS/MS

The Monarch Genomic DNA Purification kit (New England Biolabs, #T3010) was used to isolate DNA from cell pellets and eluted in LC-MS grade water. The extracted DNA was subsequently enzymatically digested into nucleosides following an established protocol ^4^. The nucleosides were injected into an Agilent 1290 Infinity UHPLC instrument with a ZORBAX Eclipse Plus C18 Rapid Resolution HD column (2.1×100 mm, 1.8 µm, Agilent #959758-902), connected to an Agilent 6495B triple quadrupole mass spectrometer operating in positive mode. The chromatographic method and mass spectrometer parameters were published previously ^4^. The data was quantified in MassHunter Quantitative Analysis for QQQ (v 10.1) using standard curves and heavy labeled internal standards for each analyzed nucleoside ^5,4^. The lower limit of quantification was 0.05 fmol for 5hmdC, 0.25 fmol for 5mdC, 0.5 fmol for dC and 1 fmol dG. The limit of detection was 0.025 fmol for 5hmdC and 5mdC as well as 0.1 fmol for dC and dG.

### Bioinformatic and statistical analysis

#### Bulk RNA sequencing data analysis

RNA-seq reads were processed using cutadapt (v4.1)^6^ to remove Illumina adapters, to quality trim at Q20, and to filter read pairs containing N bases or where either read <31 bps. Processed reads were quantified with Salmon (v1.9.0) ^7^ using transcripts from the GRCh38 Ensembl v107 annotations. Salmon’s expectation maximisation procedures were set to enable modelling of sequencing and GC biases.

Within R, transcript level counts from Salmon were imported and aggregated to the gene level using tximport v1.32.0^8^. Sample quality assessments were performed in relation to collected RNA integrity (RIN) values, sample concentrations used in library preparation (ng/ul), read GC content, and percentage read mapping. Normalisation, further PCA- and clustering-based quality control based on normalised values, and differential expression analyses were performed with DESeq2 v1.44.0^9^. Independent hypothesis weighting was conducted to optimise the power of p-value filtering using IHW v1.32.0^10^ and log2 fold change shrinkage was performed using ashr v2.2-63^11^ to reduce the impact of low expression on estimation of fold change values. Multiple correction testing adjustments were optimised by passing thresholds for both false discovery rate and log2 fold change at the point of results generation within DESeq2. Significance was assessed using a *q* value of 0.01.

### Single cell transcriptomic data analysis

#### Single cell transcriptomics data preparation

Single-cell RNA sequencing data were aligned against the probe reference file v1.0.0 from CellRanger using cellranger v7.0.0. Sample demultiplexing and read count quantification were performed with the command cellranger multi. Downstream analyses were conducted on droplets marked as non-empty according to the cutoff on the barcode rank plot of the CellRanger algorithm. Loading and processing of the data were performed with Python v3.8.10 and R v4.2.1. The count matrices were loaded using Scanpy v1.9.3 ^12^. Low-quality cells’ barcodes were identified and filtered out according to manually set thresholds, specific for each sequencing run, on the distribution of total counts and the number of genes in the droplets. Droplets with a percentage of mitochondrial genes counts higher than 5% were also discarded. Doublets were identified and removed employing scDblFinder v1.12.0 ^13^. Genes expressed in less than 150 cells were excluded from the analysis.

Count normalization was performed using the function normalise_total from Scanpy, excluding the highly expressed genes from the computation and imposing as target sum of the library 50,000 counts. Data was then log-transformed with the function log1p from the same library. Runs were subsequently integrated using Harmony, correcting for sequencing run ^14^ after computing the first 50 components of the principal component analysis (PCA).

We ultimately obtained 27925 high-quality cells: 7733 and 9571 cells derived from three samples of hEGCLCs and day 6 hPGCLCs aggregates, respectively; 6900 from two samples of hiPSCs and 3721 from one sample of iMeLCs, all of CTL08A genotype.

#### Highly variable genes selection and dimensionality reduction

For the selection of the highly variable genes, Triku v2.1.3 ^15^ was employed after a k-nearest neighbor (kNN) graph was computed on the first 11 principal components of the corrected Harmony PCA and with a number of neighbors equal to 83. A new principal component analysis was computed exclusively on the highly variable genes, then cells from each run were integrated using Harmony and a new kNN graph was computed employing the same parameters mentioned above for number of PCs for the new integrated PCA and number of neighbors. To visualize the cells of the dataset, Uniform Manifold Approximation and Projection (UMAP) algorithm was employed for the computation of a lower dimensional space ^16^, using default parameters of the UMAP function in Scanpy.

#### Clustering and annotation

For cell type annotation, cells were first clustered using the Leiden algorithm ^17^ with resolution 1. Then, clusters were annotated using a supervised approach, taking into account the expression of known gene markers (Table S2).

### Pseudobulk computation and differential gene expression analysis

#### Pseudobulk was computed by summing the gene counts from the raw counts for each replicate and cell type using the function get_pseudobulk from decoupler v1.3.4

Differentially expressed genes were investigated for the pairs hPGCLC-hiPSC, hPGCLC-hEGCLC and hiPSC-hEGCLC. For each comparison, we kept cells from samples annotated from the two cell types under investigation and genes were filtered out if not expressed in at least 100 cells. Additionally, genes were filtered out at the pseudobulk level if their number of counts was less than 10 counts in each sample or less than 20 across all samples. Normalization factors and a dispersion estimation were computed using calcNormFactors and estimateDisp from edgeR v3.40.2. Each contrast was modeled by fitting a negative binomial generalized log-linear model using glmFit on a simple model matrix taking as covariate the cell types with the formula “∼ celltype”. Differentially expressed genes (DEGs) were identified based on an FDR < 0.01, an absolute log fold change > 2, and a likelihood ratio greater than 1.

#### Longitudinal gene expression clustering

For longitudinal characterization of gene expression patterns in hiPSC, iMeLC, hPGCLC and hEGCLC, genes were clustered based on their log-fold change against a common baseline, namely hiPSC.

Briefly, pseudobulk profiles were derived for each sample from cells annotated as one of these four cell types, as described above. Genes were filtered if not expressed in any sample, in at least 100 cells or in at least 10% of the cells of a sample. Additionally, genes were filtered out if they had less than 10 counts per sample or less than 200 total counts across samples. Counts were normalized and an estimation of the dispersion was performed as above. The model matrix was designed with the formula “∼ celltype” and reference “hiPSC”. The negative binomial generalized log-linear model was performed using glmFit and returned the log-fold change for each cell type against the hiPSC.

For subsequent clustering, genes of the differential testing were kept only if their FDR < 0.01 and if the absolute log-fold change was higher than 1 in at least one cell type. The log fold-changes were standardized removing the mean and scaling to unit variance. The clustering task was performed using an agglomerative approach using a Euclidean linkage distance, the “ward” linkage criterion and setting 15 as the total number of clusters.

#### Functional enrichment analysis

Functional enrichment analysis on differentially expressed gene sets and the clusters of genes defined by the longitudinal clustering approach was performed by using topGO^18^ (R package version 2.52.0.), to identify significant overlaps with Gene Ontology terms’ gene sets, and decoupleR^19^, for enrichment analysis of KEGG and Reactome terms. After the definition of the gene set of interest, Fisher’s exact test was employed to define the significance of the overlap. When using topGO, the weight01 algorithm was employed to account for the hierarchical structure of the ontology. Terms were excluded if the number of annotated genes of the term was less than 15 and more than 200 or if the number of significant genes found in the gene set was lower than 5. An enrichment score was calculated for each term as the ratio of the number of significant genes and the number of expected ones. A term was deemed significant if its p-value was lower than 0.01 and its enrichment score was higher than 2.

#### Chen, et al., in vitro external dataset preprocessing

Raw count matrices from Chen et al. ^20^ were downloaded from GEO database from the series with accession code GSE140021, specifically runs including batch 1 runs from the hESC cell line with high competency named UCLA2, that is also the male line (accession codes from GSM4202941 to GSM4202946). Cells were filtered as in the original publication, keeping those expressing at least 200 genes and a maximum of 8000 genes. Additionally, cells having a percentage of mitochondrial genes higher than 20% or a percentage of ribosomal genes higher than 40% were discarded. Only genes present in at least three cells were considered in the analysis. The UMI count for each cell was normalized by total expression, scaled by multiplying by 10000 and then log-transformed. The top 2000 highly variable genes were selected with the Scanpy function “highly_variable_genes” and were used to compute the principal component analysis. A k-nearest neighborhood was computed on the first 50 principal components using 20 as the number of nearest neighbors. After a first clustering with Leiden algorithm at resolution 0.8, three clusters were excluded because, upon close inspection, they revealed poor quality. The neighborhood graph was recomputed on the first 9 principal components and a new Leiden clustering was performed. We annotated the clusters based on the expression of known markers (Table S2).

#### Guo et al., and Chitiashvili et al., ex vivo external datasets preprocessing

For Guo et al., ^21^ data of single-cell RNA-seq for embryonic and fetal human testes were downloaded from GEO (accession code GSE143356). These were jointly analysed with data of single-cell RNA-seq for embryonic and fetal human ovaries from Chitiashvili et al. (^22^ accession code GSE143380). First, all libraries were merged, and cells were subjected to the same filtering: cells were excluded if expressing less than 250 genes or more than 8000 and if having less than 350 total counts or more than 50000. Additionally, cells having a percentage of mitochondrial genes higher than 10% or a percentage of ribosomal genes higher than 50% were excluded. Genes were excluded from the analysis if expressed in less than 5 cells and if having zero counts. The UMI counts for each cell were normalized based on the total expression using the default parameters of normalize_total, followed by log transformation. Highly variable genes were identified using Scanpy’s default method, and the data were scaled to account for variation due to UMI counts and mitochondrial gene expression. Principal component analysis was computed and the k-nearest neighbourhood was computed with default parameters of the “neighbours” function from Scanpy. Cells were clustered by Louvain algorithm with resolution = 0.5 and the UMAP package was used to visualize cells. Annotation was performed by expression of the markers indicated in the original publications ^21,22^.

#### Ingestion on external datasets

The single-cell transcriptomic data produced in this study were projected separately onto the principal component analysis space of two external datasets (one in vitro and another one of in vivo fetal data), using the ingest function from Scanpy. For both ingestions, we subset the genes of the external datasets on the genes in our data and computed a new principal component analysis onto which we built a new kNN graph and visualization using the same number of principal components and neighbours as the one specified in the preprocessing of each dataset.

#### Transcription factor activity analysis

To infer transcription factors’ (TFs) activities in our data, a Multivariate Linear Model method was employed on the unnormalized gene expression using the decoupler implementation. The regulatory adjacency matrix used as covariate of the model was downloaded from the Dorothea database through decoupler’s API^19,23^ and only the evidence level A, B and C were kept in the matrix, resulting in 292 TFs and their 14582 targets. The estimated t-values were taken into account as the TFs’ activities, as suggested by the tool. Pearson correlation coefficient was calculated between two cell types based on the mean activity values of each transcription factor.

#### DNA methylation data analysis

Data were preprocessed using nf-core/methylseq v2.3.0 pipeline (https://nf-co.re/methylseq/2.3.0) ^24^, built in Nextflow v22.10.1 ^25^. The quality of FastQ files was evaluated with FastQC (https://www.bioinformatics.babraham.ac.uk/projects/fastqc/) v0.11.9. Reads were then trimmed by TrimGalore! (https://www.bioinformatics.babraham.ac.uk/projects/trim_galore/) v0.6.7 with default parameters for the removal of adapter contamination and low-quality regions. Furthermore, 8 additional bases from the ends of both R1 and R2 reads were trimmed. bwa-meth (https://github.com/brentp/bwa-meth) ^26^ v0.2.2 was used to align trimmed reads to GRCh38.p13 (hg38) reference genome, applying MarkDuplicates from Picard (http://broadinstitute.github.io/picard) v2.27.4 to remove duplicated reads. Additionally, we calculated quality metrics specific for targeted sequencing experiments, e.g., off-bait percentage, using Picard’s CollectHsMetrics function (COVERAGE_CAP = 1000; NEAR_DISTANCE = 500, default values for remaining parameters). MethylDackel (https://github.com/dpryan79/MethylDackel) v0.6.1 was then used for the extraction of cytosine methylation calls. The methylation values from the two strands were added to obtain a single value describing the double-strand CpG methylation level using MethylDackel’s mergeContext function. MultiQC v1.13 ^2727^ was used to summarize QC results. Finally, we filtered bedGraph files keeping only on-bait CpG loci covered by at least 10 reads.

All the following analyses were done in R v4.2.1. CpG loci from all examined samples were collected into a single BSseq object using bsseq ^28^ v1.34.0 R package. Only CpGs called in at least 75% of the samples were kept using MethylSig ^29^ v1.10.0 R package. Two group comparisons to find differentially methylated regions (DMRs) were performed with DSS ^30^ v2.46.0 R package. Smoothing was applied in estimating mean methylation levels (by cell type). DMRs were selected setting delta > 0.25 and p.threshold < 0.0001, while the remaining parameters were left as default. DMRs were annotated with ChIPseeker ^31,32^ v1.34.1 R package, which associated DMRs with genes whose transcription start sites (TSSs) are closest to the center of DMRs (default parameters, except TxDb object). Annotation with genomic region types was also performed by considering promoter regions located ±3000 bp from TSS. The TxDb object containing transcript-related features was given in input, after creating it from the GTF file of the GENCODE Human Release 35 (Ensembl v101) (https://www.gencodegenes.org/human/release_35lift37.html). Gene annotation was performed with biomaRt ^33^ v2.54.1 R package, selecting as host https://aug2020.archive.ensembl.org. DMRs were visualized with the plotDMRs function of the dmrseq ^34^ v1.18.1 R package, which also exploits annotatr ^35^ v1.24.0 R package to retrieve the CpG annotation visible in the plot. The sechm (doi: https://doi.org/10.18129/B9.bioc.sechm) v1.6.0 R package was used for heatmaps. Functional enrichment analysis of the differentially methylated genes (DMGs), i.e. genes annotated to the DMRs, was done as previously described in the case of scRNAseq analysis, except for considering as universe gene set for the enrichment only the genes covered by the Twist baits. The explored imprinted regions with their annotated genes were taken from ^36–38^. These regions were overlapped with the BSseq object to explore their methylation levels in our samples. The explored escapee regions were the ones found in ^37^. Those covered by at least 10 CpG in all ^37^ human primordial germ cell (hPGC) samples and with methylation levels above 30% in ^37^ week 5-9 hPGC samples were selected. The conversion of genomic coordinates from assembly GRCh37 (hg19) to GRCh38 (hg38) was performed. As for imprinted regions, also escapee regions were overlapped with the BSseq object for exploratory purposes and to retrieve the so-called escapee CpGs, i.e., single CpGs within escapee regions. The latter were annotated as previously described for the DMRs. Functional enrichment analysis was made using the genes annotated to the escapee CpGs, following the same approach described above. The escapee CpGs were also compared with non-escapee CpGs, defined as all other CpGs in the BSseq object, that were not within the defined escapee regions. The distribution of methylation levels of the 9113 escapee CpGs was tested against that of 500 sets of 9113 CpGs randomly selected within the non-escapee regions.

#### Directed gene regulatory networks deconvolution

A set of cell type-specific regulatory regions was generated by integrating data from previously published ATAC-seq data from hESC, iMeLC and hPGCLC (GSE 120648, ^39^). Published bigwig files were filtered with a threshold of 5 normalized counts, and consecutive covered bins were merged to form individual open peaks. Enhancers were identified as accessible regions spanning between 100 and 1,000 bp, and associated to the 2 closest promoters within 50,000 bp. Only promoters and enhancers of genes expressed in each cell type were considered for the following analyses.

Gene regulatory networks (GRNs) were built using CellOracle (^40^, v. 0.10.12). All peaks in the previously generated list of accessible promoters and enhancers were scanned for the presence of TF motifs from the gimmemotifs database (gimme.vertebrate.v5.0) using gimme motif scan through the CellOracle pipeline. Base GRNs were then generated for each combination of cell types, connecting each TF to the genes (among the HVGs and selected genes, listed in Table S4, to ensure that we include all genes expressed in our scRNAseq dataset) whose enhancers or promoters bore the TF binding motif. Through CellOracle we leveraged co-expression patterns to generate GRNs based on cell type-specific gene expression derived from scRNAseq data from this paper. Edges in the resulting networks are drawn based on the concurrence of the following criteria: presence of the source TF’s binding motif in the gene promoter or its enhancers, co-expression of the TF and the gene, and the discernible impact of TF expression on the transcript levels of the gene, inferred by a Bagging Ridge ML model. The GRNs were then processed using custom bash/R scripts and visualized with Cytoscape (^41^, v.3.10.0), selecting the yFiles Orthogonal layout algorithm.

#### Master regulators analysis

Master regulators analysis (MRA) was performed to identify TFs in the GRNs (built as described above) significantly involved in dMdE genes regulation, filtering those with p-value < 0.05 and enrichment>2. dMdE (differentiallyMethylated-differentiallyExpressed) genes were selected as follow: genes belonging to pseudo-bulk clusters 2 and 6 (i.e upregulated in hPGCLC) intersected with target genes of regulatory regions significantly less methylated in hPGCLC when compared to both hiPSC and hEGCLC were united with the ones belonging to pseudo-bulk cluster 1 and 5 (i.e downregulated in hPGCLC) intersected with targets of regulatory regions significantly more methylated in hPGCLC when compared to both hiPSC and hEGCLC. To calculate enrichment, we compared the percentage of targets of each TF in the dMdE group with the percentage of targets of the same TF among the expressed genes. To calculate the p-value of each enrichment we performed a hypergeometric test overlapping dMdE genes that were targets of a certain TF with all dMdE genes, using as background expressed genes in the single-cell RNA-seq data.

## Data availability

Bulk RNA-seq, single-cell RNA-seq and DNA methylation data generated in this study will be available at NCBI Gene Expression Omnibus and Sequence Read Archive.

## Code availability

Full code used for the analyses will be available at https://github.com/GiuseppeTestaLab/EGCLC_paper_release

## Supplementary figures

**Figure S1.**
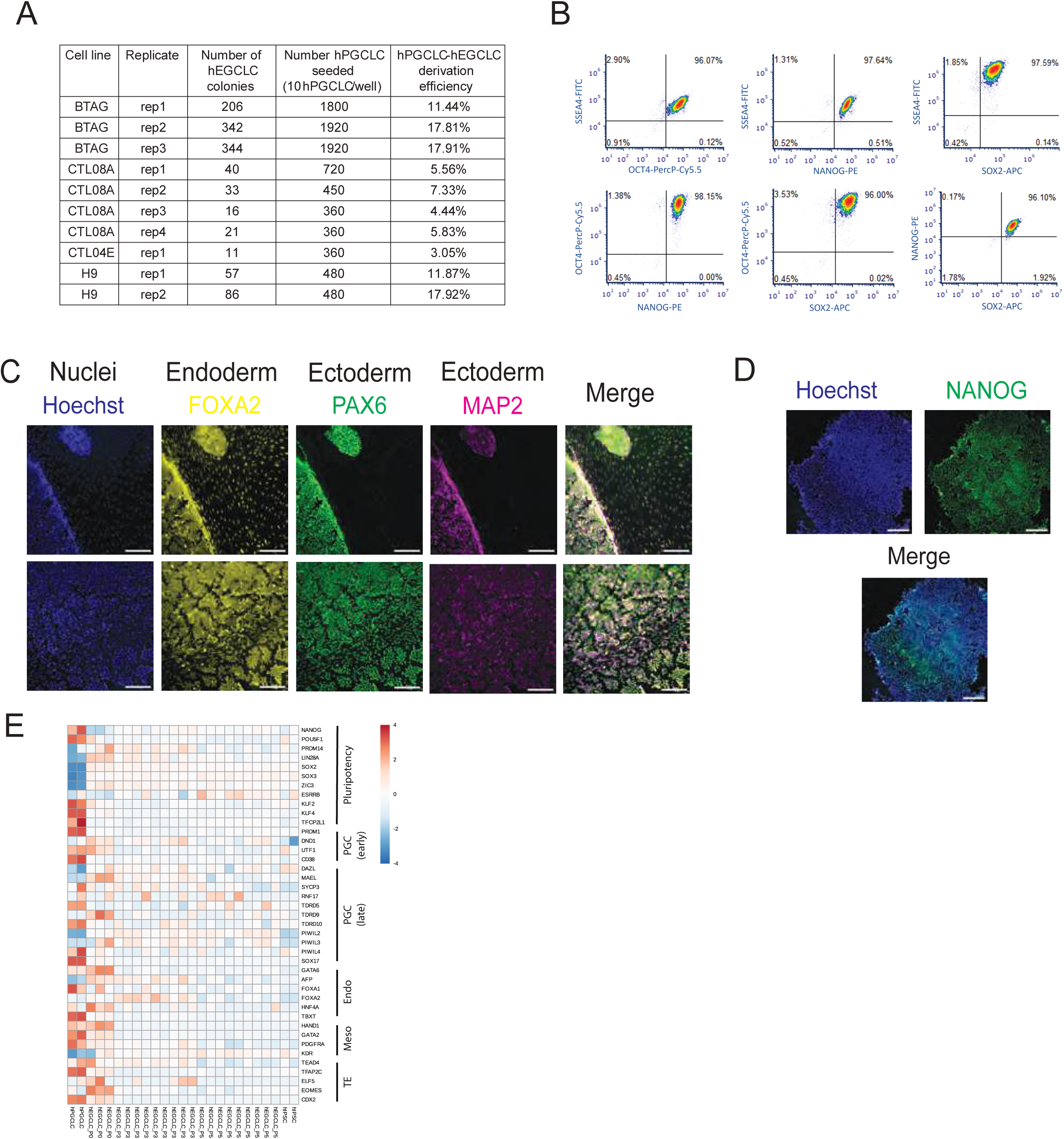
hEGCLC pluripotency validation, related to Figure 1. **(A)** Table summarizing hEGCLC reprogramming efficiency (calculated as total number of hEGCLC colonies, obtained 14 days after hPGCLC seeding, divided by total number of hPGCLC seeded [ie. 10 x total number of well seeded, since we plated 10 hPGCLC/well]) for each of the hPSC lines used. **(B)** Representative flow cytometry density plots of pairwise antibody combinations for pluripotency markers (SOX2, OCT4, NANOG and SSEA4) of hEGCLC; percentages of positive cells are shown. **(C)** Representative images of hEGCLC colonies immunostained for pluripotency markers: SOX2, OCT4; nuclei stained with Hoechst (scale bar: 250 μm) **(D)** hEGCLCs tri-germ layer differentiation: immunostaining for lineage specification markers: FOXA2 (endoderm), PAX6 and MAP2 (ectoderm); nuclei stained with Hoechst (scale bar: 150 μm) **(E)** Heatmap of bulk expression of relevant genes (rows), grouped in the following categories: pluripotency, PGC (early), PGC (late), Endoderm, Mesoderm and Trophoectoderm (TE) markers. Samples are shown in columns (replicates of day4 hPGCLC, hEGCLC P0, P3 and P5 and hiPSC). Color gradient indicates Z-score.

**Figure S2.**
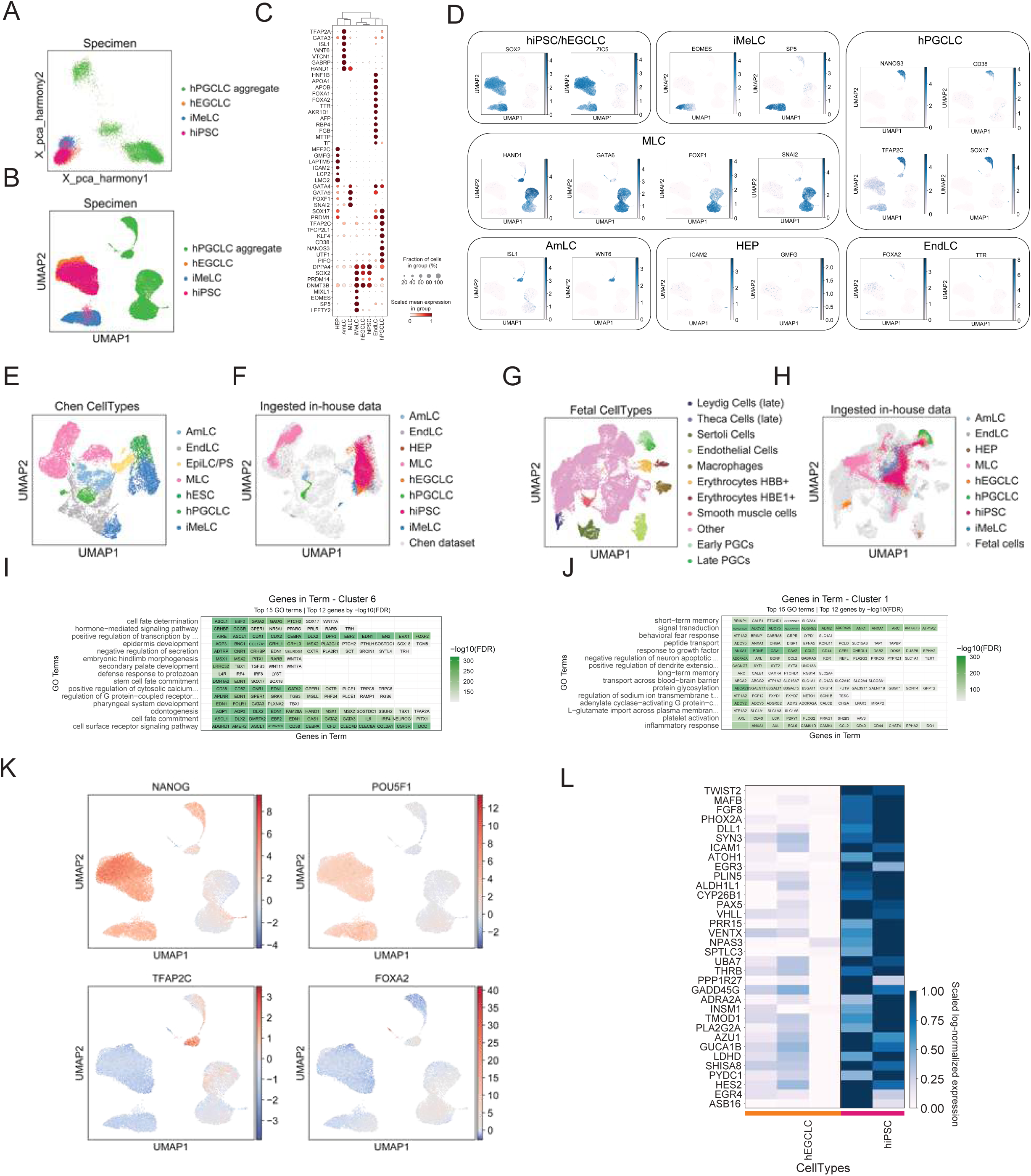
In vitro model single-cell transcriptomic profiling and benchmarking onto in vitro and in vivo fetal references, related to Figure 2 **(A)** PCA of the samples indicated in the figure legend (day 6 hPGCLC aggregates; hiPSCs = human induced Pluripotent Stem Cell; iMeLCs = incipient Mesoderm-Like Cells; hEGCLCs = human Embryonic Germ Cell-Like Cells). **(B)** UMAP of all cells in the dataset, after pre-processing and filtering, colored by samples metadata (n = 2 samples for hiPSCs, n = 1 sample for iMeLCs, n = 3 samples for day6 hPGCLC aggregates and hEGCLCs) **(C)** Dot plot of gene expression for some of the relevant markers used in the annotation. Dot size is proportional to the number of cells expressing that marker and color encodes the mean expression in group (log norm counts). **(D)** UMAP of all cells in the dataset colored by expression of relevant markers used in the annotation. Plotted markers are divided by the cell type they are most relevant for. **(E)** UMAP of the UCLA2 external dataset (Chen et al. 2019), after pre-processing and filtering, colored by annotated cell types (hESC = human Embryonic Stem Cell; iMeLC = incipient Mesoderm-Like Cells; hPGCLC = human Primordial Germ Cell-Like Cells; MLC = Mesoderm-Like Cells; AmLC = Amnion-like cells; EndLC = Endoderm-like cells; EpiLC/PS = Epiblast-like Cell/Primitive Streak). **(F)** Projection of our scRNA-seq data (colored according to cluster annotation) on the *in vitro* external reference dataset (Chen et al. 2019), in grey. **(G)** UMAP of the external datasets(Guo et al. 2021; Chitiashvili et al. 2020) (human prenatal gonads [6-16 weeks post-fertilization (wpf)]), after integration, pre-processing and filtering, colored by annotated cell type (PGC = primordial germ cell). **(H)** Projection of our scRNA-seq data (colored according to cluster annotation) on *in vivo* fetal datasets (Guo et al. 2021; Chitiashvili et al. 2020) (in grey). **(I)** and **(L).** Top 12 genes among the differentially expressed genes (FDR < 0.01 and absolute logFC > 2), ordered by false discovery rate (FDR) for each of the top 15 GO terms shown in Figure 2E (cluster 6, upregulated in hPGCLC) and 2F (cluster 1, downregulated in hPGCLC), respectively. Color gradient indicates -log10(FDR). **(M)** UMAPs of our dataset colored by inferred activity of the following TFs: NANOG, POU5F1, TFAP2C and FOXA2. **(N)** Heatmap of pseudo-bulk expression of the 34 genes downregulated in hEGCLC compared to hiPSC.

**Figure S3.**
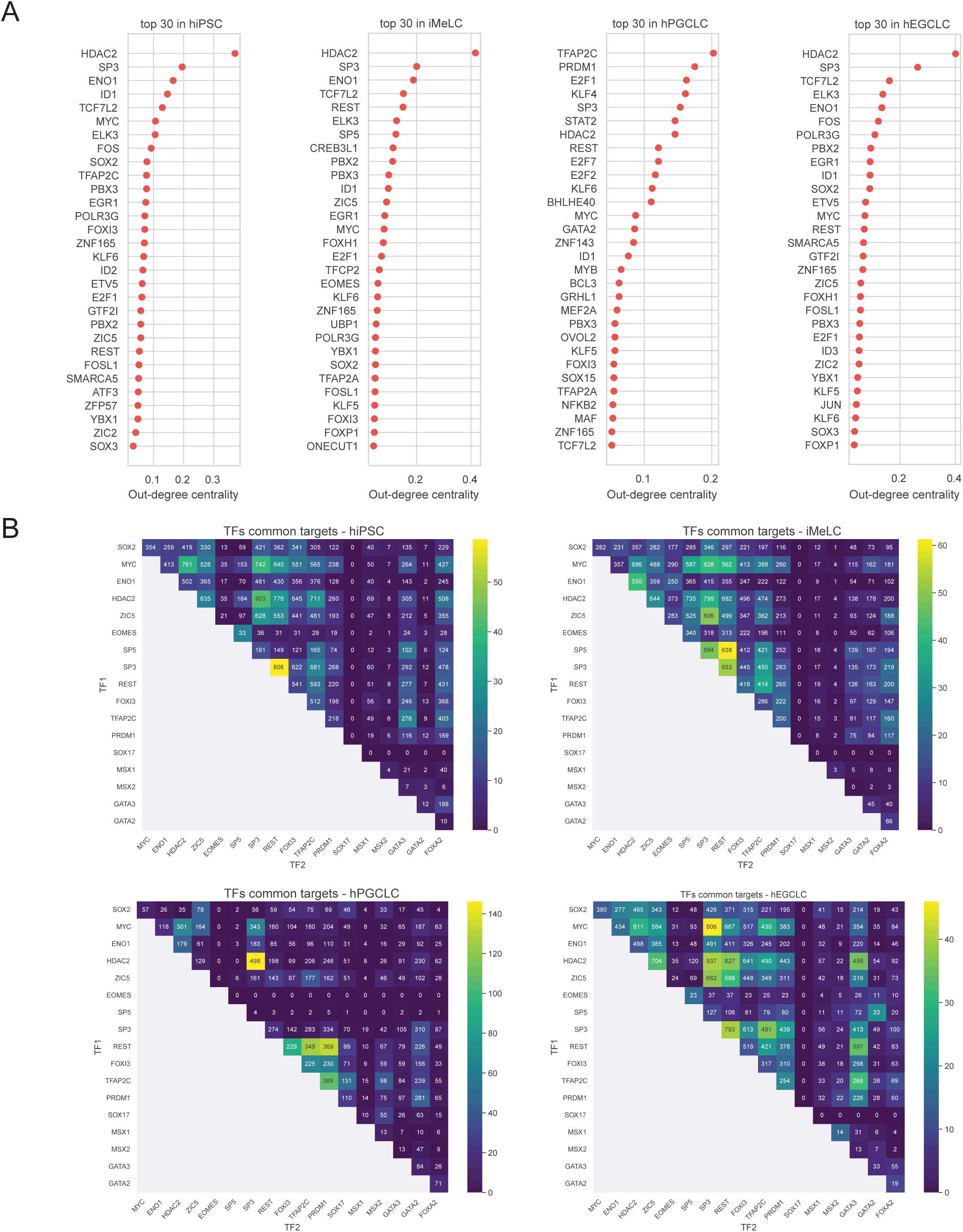
Key regulatory TFs in hiPSC, iMeLC, hPGCLC and hEGCLC GRNs, related to Figure 3 **(A)** Dot plot of top-ranking transcription factors (TF) based on degree out-centrality (X-axis) measured for each cell type (reported in each plot title). **(B)** Heatmap of TFs cooperativity for each cell type GRN (indicated in the plot title). X and Y-axis TF list. Color gradient: -log10(p-value), (p-value < 0.05). in each square is indicated the number of common targets co-regulated by the two cooperating TFs

**Figure S4.**
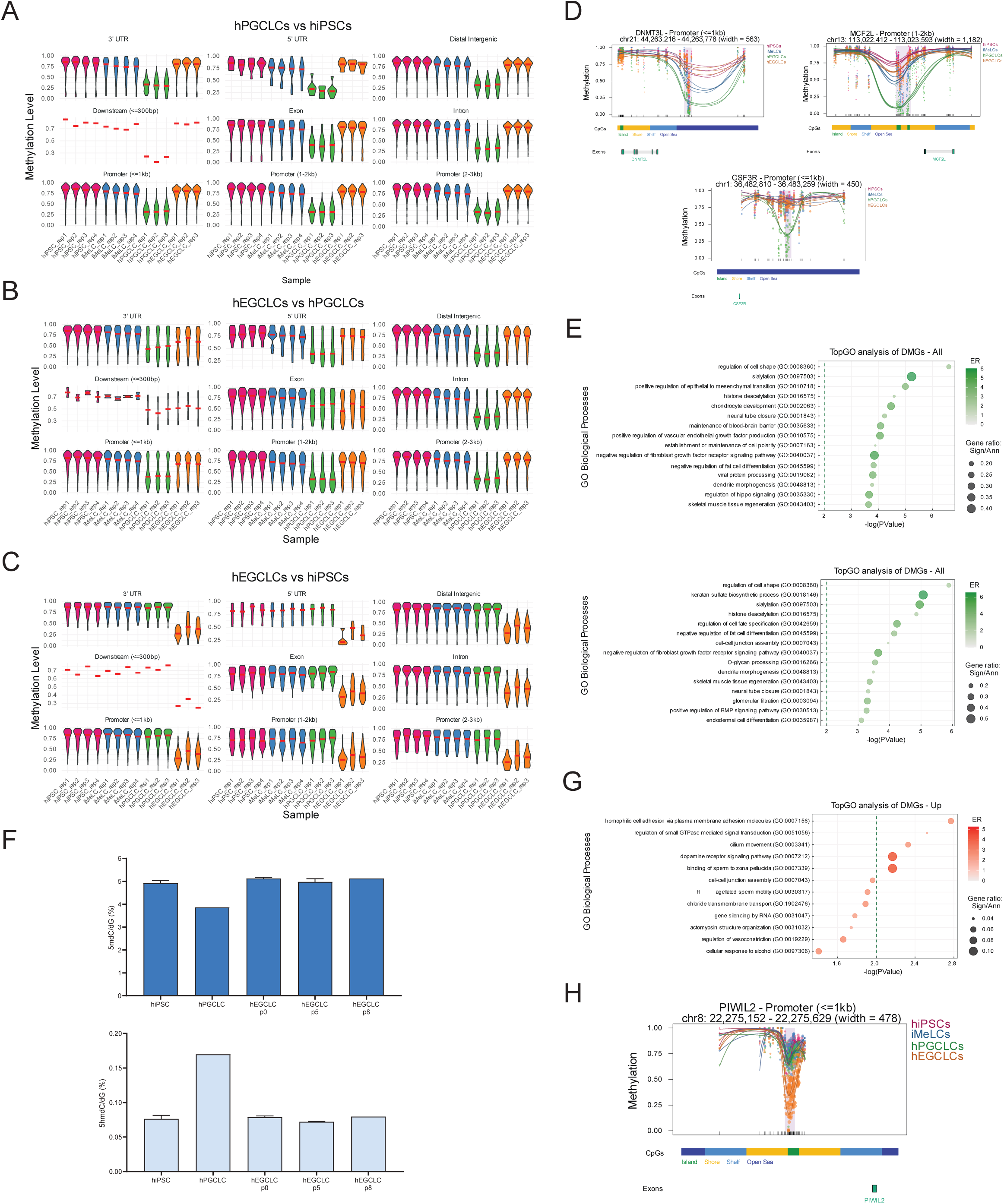
In vitro model longitudinal DNA methylation profiling, related to Figure 4 **(A), (B), (C)** Violin plots, divided by annotated genomics regions, of CpG average methylation level for each DMR (%, Y-axis) in each sample, X-axis. Red horizontal bars indicate the median methylation level. **(A)** hPGCLCs vs hiPSCs comparison; **(B)** hEGCLCs vs hPGCLCs comparison; **(C)** hiPSCs vs hEGCLCs comparison. **(D)** DMR plots of three regions showing a significant loss of methylation in hPGCLCs compared to hiPSCs and hEGCLCs. **(E)** Dot plot of enriched biological process (BP) gene ontology (GO) terms for genes associated to DMRs between hiPSCs and hPGCLCs (top panel), hEGCLCs and hPGCLC (bottom panel). Y-axis: GO term; X-axis: -log(p-value). Color gradient indicates the enrichment ratio (ER, number of observed significant differentially methylated genes divided by the number of expected genes annotated as associated to that term); dot dimension indicates gene ratio (number of observed significant differentially methylated genes divided by the number of genes annotated as associated to that term). The dotted vertical line indicates the p-value threshold at 0.01. **(F)** Bar plot showing the percentage of methylated (5mdC/dG, top panel) and of hydroxy-methylated (5hmdC/dG, bottom panel) cytosines for the samples indicated on the X-axis (BTAG hiPSC, BTAG day 4 hPGCLC, and hEGCLC p0 and p5 from BTAG and H9 cell lines) **(G)** Dot plot of enriched biological process (BP) gene ontology (GO) terms for genes associated to DMRs where methylation is higher in hiPSCs vs hEGCLCs. Y-axis: GO term; X-axis: -log(p-value. Color gradient indicates the enrichment ratio (ER>2); dot dimension indicates gene ratio. The dotted vertical line indicates the p-value threshold at 0.01. **(H)** DMR plot of PIWIL2 promoter region.

**Figure S5.**
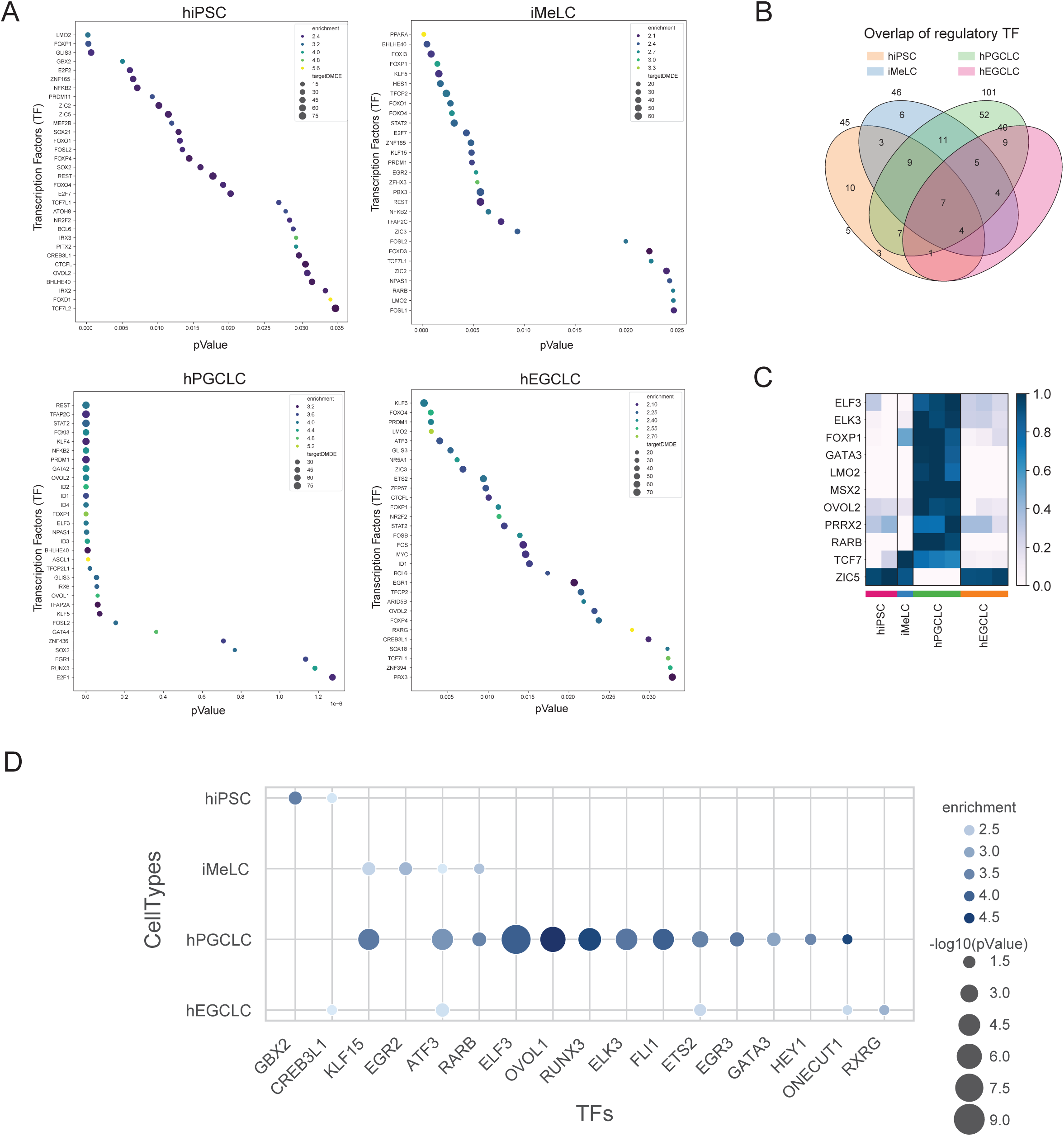
Master regulators of dMdE genes for each of the four GRNs, related to Figure 5 **(A)** Dot plot of TFs (Y-axis) significantly regulating dMdE genes, in each of the four GRNs (cell type indicated in each plot title), ranked by p-value (>0.05). Dot color: enrichment (>2); dot size: number of target dMdE genes regulated by that TF. **(B)** Venn diagram showing the overlap of TFs regulating dMdE genes among the four key cell types: hiPSC, iMeLC, hPGCLC, hEGCLC. **(C)** Heatmap of expression level (pseudo-bulk) of the 11 TFs present in the GRNs that are differentially methylated and differentially expressed and that significantly regulates dMdE in hiPSC, iMeLC, hPGCLC and hEGCLC respectively. **(D)** Dot plot of methylation sensitive TFs (X-axis) significantly regulating dMdE genes in each cell type (Y-axis), (enrichment>2, p-value< 0.05). Dot color represents enrichment and dot size represents -log10(p-value).

**Figure S6.**
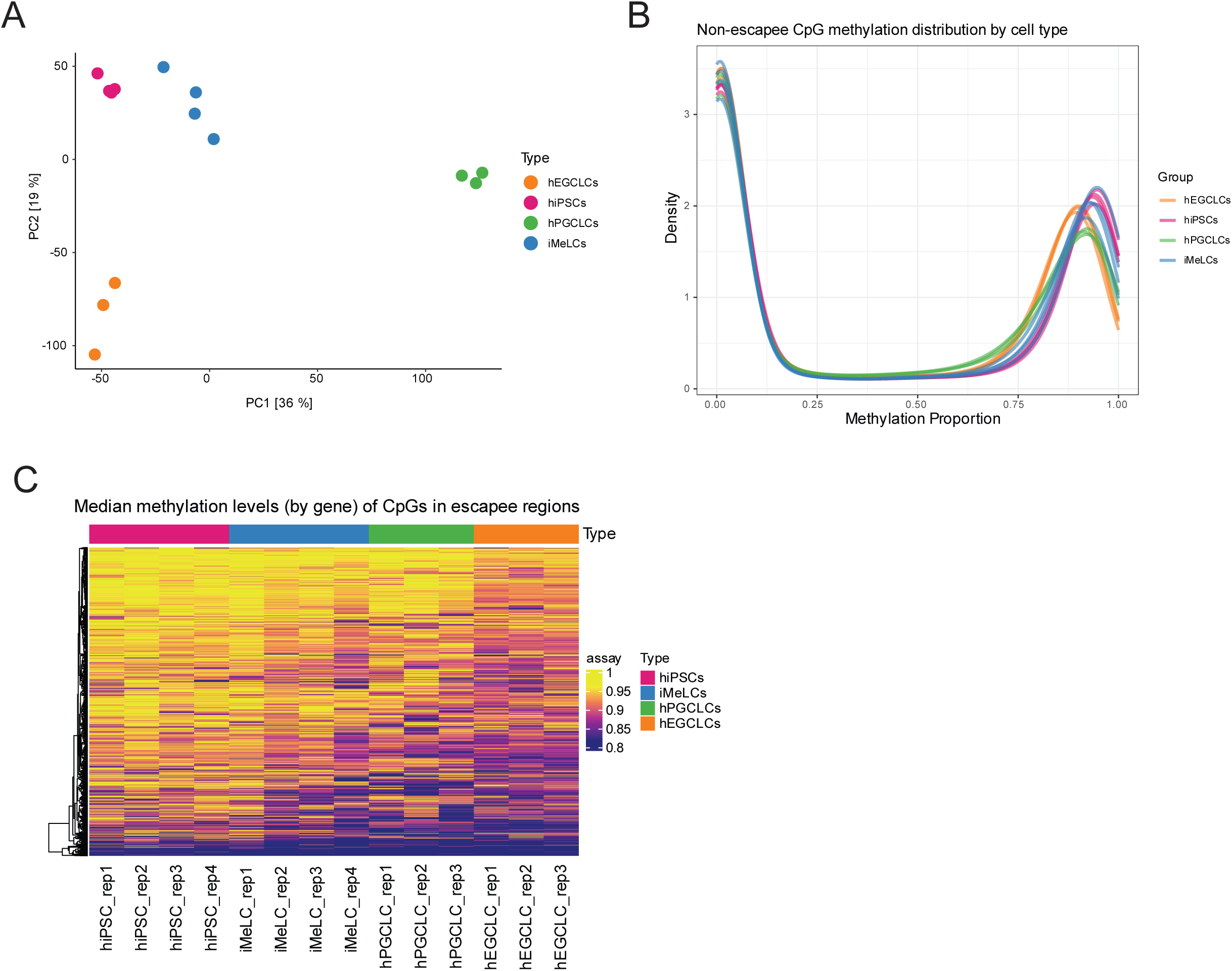
CpG methylation levels of escapee and non-escapee regions, related to Figure 6 **(A)** Principal component analysis (PCA) plot of CpG methylation levels of all non-escapee regions in hiPSCs, iMeLCs, day 6 hPGCLCs and hEGCLCs (each cell type at least in triplicate). PC1 and PC2 explain 36% and 19% of the variance, respectively. **(B)** Density plot of the distribution of CpG mean methylation levels in all non-escapee regions, by cell type **(C)** Heatmap of median methylation levels of CpGs within escapee regions for all the samples.

## Supplementary Video - Related to Figure 1

**Video S1** Time-lapse imaging of day 2 plated hPGCLC, BTAG reporter line (expression of BLIMP1-tdTomato and TFAP2C-eGFP)

**Video S2** Time-lapse imaging of day9-13 plated hPGCLC, BTAG reporter line (downregulation of BLIMP1-tdTomato and TFAP2C-eGFP)

**Video S3** Time-lapse imaging of day4-8 plated hPGCLC - SOX2 reporter line (upregulation of SOX2-dTomato)

## List of Supplementary Tables

**Table S1** - bulkRNAseq differentially expressed genes List of bulkRNAseq differentially expressed genes (DEGs) for the following comparisons (BTAG cell line), related to Figure 1G, 1H: A. hiPSC-hEGCLC passage 0 B. hiPSC-hEGCLC passage 3 C. hiPSC-hEGCLC passage 3

**Table S2** - Markers used to annotate scRNAseq datasets Markers used to annotate cell type of the following scRNAseq datasets, Related to Figure 2 and S2: A. scRNAseq dataset from this paper (on CTL08A line) B. scRNAseq dataset from Chen et al., 2019 C. scRNAseq fetal dataset from Guo et al., 2021 and Chitiashvili et al., 2020 D. Distributions of annotated cell types in Leiden clusters

**Table S3** - scRNAseq differentially expressed genes and related Gene Ontology terms Gene list for each of the 15 clusters obtained from longitudinal gene expression clustering of CTL08A scRNAseq (pseudobulk) and for hiPSC-hEGCLC DEGs and Gene Ontology terms for each of these gene lists, related to Figure 2 and S2 A. Gene list for each of the 15 clusters B. Go terms associated to the gene lists of each of the 15 clusters C. hiPSC-hEGLC DEGs (pseudobulk) D. Go terms associated to genes upregulated in hEGCLC vs hiPSC

**Table S4** - Gene regulatory networks (GRN) for each cell type List of selected genes added during GRNs generations and cell type specific GRNs (source TF and target gene pairs and associated GRN parameters), related to Figure 3 and S3 A. hiPSC GRN B. iMeLC GRN C. hPGCLC GRN D. hEGCLC GRN E. list of selected genes added during GRNs generation, to ensure the inclusion of all genes expressed in our scRNAseq dataset

**Table S5 -** Differentially methylated regions and related Gene Ontology terms List of differentially methylated regions (DMRs) for each of the comparisons listed below and related Gene Ontology terms, relative to Figure 4 and S4. A. Median CpG methylation levels for each cell types B. hiPSC-iMeLC DMRs and Go terms C. hiPSC-hPGCLC DMRs and Go terms D. hiPSC-hEGCLC DMRs and Go terms E. iMeLC-hPGCLC DMRs and Go terms F. iMeLC-hEGCLC DMRs and Go terms G. hPGCLC-hEGCLC DMRs and Go terms

**Table S6** - Imprinted regions A. List of imprinted regions, relative to Figure 4H

**Table S7** - Methylation sensitive transcription factors List of methylation sensitive TFs (from Yin et al., 2017) that we intersected with our list of TFs, related to Figure S5D

